# A mutation-led search for novel functional domains in MeCP2

**DOI:** 10.1101/288878

**Authors:** Jacky Guy, Beatrice Alexander-Howden, Laura FitzPatrick, Dina DeSousa, Martha V. Koerner, Jim Selfridge, Adrian Bird

**Author notes:** Corresponding Authors. JG, AB.

## Abstract

Most missense mutations causing Rett syndrome affect domains of MeCP2 that have been shown to either bind methylated DNA or interact with a transcriptional co-repressor complex. Several mutations, however, including the C-terminal truncations that account for ~10% of cases, fall outside these characterised domains. We studied the molecular consequences of four of these “non-canonical” mutations in cultured neurons and mice to see if they reveal additional essential domains without affecting known properties of MeCP2. The results show that the mutations partially or strongly deplete the protein and also in some cases interfere with co-repressor recruitment. These mutations therefore impact the activity of known functional domains and do not invoke new molecular causes of Rett syndrome. The finding that a stable C-terminal truncation does not compromise MeCP2 function raises the possibility that small molecules which stabilise these mutant proteins may be of therapeutic value.

## Introduction

Mutations in the MECP2 gene cause the profound neurological disorder Rett syndrome (1) (RTT, OMIM #312750). The gene product, MeCP2 protein, is expressed in most cell types, but is especially abundant in neurons (2–4). Accordingly, loss of MeCP2 function has the most profound effect in the nervous system, with minimal phenotypic consequences in other tissues (3, 5, 6). Because of its association with intellectual disability in RTT and other disorders, the MECP2 gene is frequently screened for mutations in clinical cases of developmental delay. This has defined many amino acid changes as RTT-causing or as relatively benign or neutral variants that do not cause Rett syndrome (7, 8). The MeCP2 protein sequence is 95% identical between rodents and humans, suggesting stringent purifying selection throughout. Nevertheless, RTT missense mutations are non-randomly distributed in two prominent clusters, whereas apparently neutral polymorphisms are broadly distributed across the remainder of the protein (Fig. 1). The implication that RTT mutational clusters mark critical domains is supported by biochemical studies, which show that mutations within the methyl-CpG binding domain (MBD) disrupt DNA binding, whereas mutations within the NCoR Interaction Domain (NID) prevent binding to the NCoR/SMRT corepressor complexes (9–11). High-resolution X-ray crystallographic structures of both these interactions support the view that MeCP2 bridges methylated sites on DNA with a histone deacetylase-containing corepressor (12, 13). Further evidence for the importance of these two domains comes from a recent study in which mice carried a pared down MeCP2 “minigene”, ΔNIC, consisting of only the MBD, a nuclear localisation signal and the NID (14). These mice survived for over a year with only mild symptoms. A simple interpretation is that this interaction restrains transcription; a view supported by recent transcriptomic analyses of MeCP2-depleted mouse brain (15–17).

**Figure 1.**
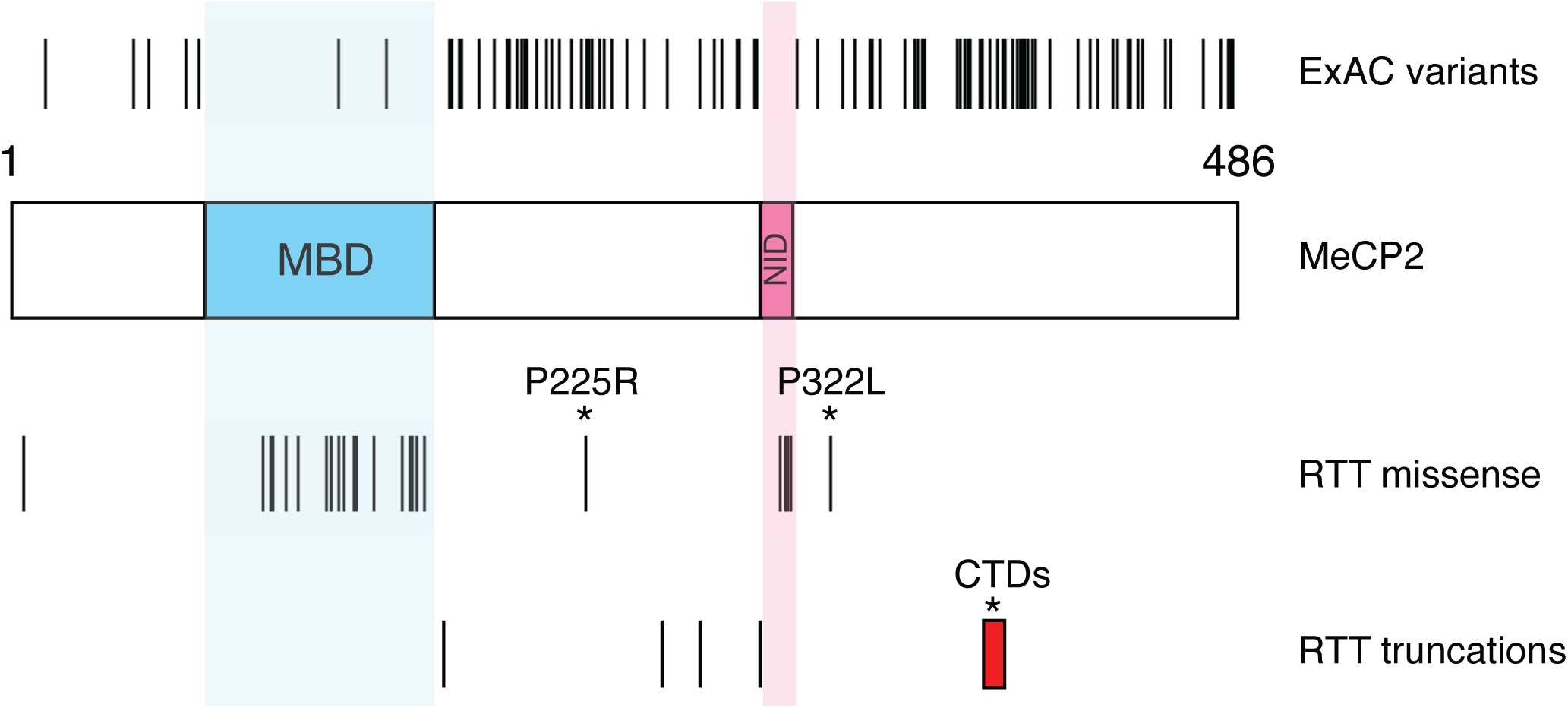
Population variants and RTT mutations in the coding sequence of MeCP2. A schematic diagram of human MeCP2 protein shows the methyl-binding domain (MBD) and NCoR/SMRT interaction domain (NID). Single amino acid changes found in males in the ExAC exome sequence database are indicated above the protein and RTT-causing mutations are shown below. Only male ExAC changes are shown to exclude the possibility that pathogenic mutations in females could be masked by skewed X-inactivation. All RTT mutations have been shown to be absent in the parents in at least one case. Red box – range of deletion start points for C-terminal deletion (CTD) RTT frameshift mutations. The “non-canonical” RTT mutations that are the subject of this study are marked with asterisks.

A significant limitation of this model for MeCP2 function is that several recurrent mutations, whose causal role in RTT is confirmed by their absence in both parents, lie outside the MBD and NID (Fig. 1) and are absent in Δ NIC mice (14). Most prominently, about 10% of all RTT cases involve a heterogeneous set of small deletions in exon 4 of *Mecp2.* The resulting frameshifts introduce premature translational termination often preceded by stretches of novel amino acid sequence of variable length (18, 19). As these C-Terminal Deletions (CTDs) are far downstream of the MBD and NID, they hint that there are other, as yet undetected, functional domains within MeCP2. The same argument applies to two other recurrent RTT mutations, P225R and P322L, which lie outside the minimal regions required for DNA and NCoR binding. Here we use biochemical, cell biological and genetic methods to show that these mutations recapitulate RTT-like phenotypes in mice through either or both of two mechanisms: 1) by destabilising MeCP2, so that only a fraction of the wild-type level remains; 2) by interfering with the ability of MeCP2 to recruit the NCoR complex to chromatin and repress transcription. Importantly, we find that one C-terminal truncation is stable and functionally wild-type (WT) in a mouse sequence context, but unstable and RTT-causing when the protein is “humanised”. This demonstrates that the truncation itself only has a measurable functional consequence if it leads to destabilisation. From this and other data we conclude that these “non-canonical” RTT mutations do not identify previously unappreciated functional domains. Moreover, our results highlight the crucial role played by the MeCP2 protein sequence in ensuring stability.

## Results

### P225R and P322L mutations cause Rett-like phenotypes in mice

To test the hypothesis that the MBD and NID are key to MeCP2 function, we first identified RTT mutations that apparently leave these domains intact. Next, we generated mouse models to check that these recapitulated RTT-like phenotypes. Finally, we used cellular and molecular assays to ask whether these mutations affect the functions that have been attributed to the MBD and the NID. According to our rationale, new functional domains would be expected to cause RTT without compromising either binding to DNA or NCoR recruitment.

We initially selected the P225R and P322L mutations, which occur in a relatively small number of Rett syndrome cases (Table 1). Parental analysis has shown that these missense mutations arise *de novo* in the affected individuals (7), strongly suggesting that they are causative. We modeled these mutations by gene targeting in mouse ES cells (Fig. 2a; Supp. Fig. 1a) to produce mouse alleles that differed from WT only at the patient mutation site and a single loxP site remaining in intron 2 after deletion of the selection cassette (Fig. 2a; Supp. Fig. 1b-e). To determine whether P225R and P322L cause RTT-like phenotypes in mice, we injected mutant ES cells into blastocysts to create chimeras and bred them to establish *Mecp2^P225R^* (P225R) and *Mecp2^P322L^* (P322L) mouse lines (Supp. Fig. 2a,b). P322L hemizygous males appeared normal at weaning, but from approximately 5 weeks of age they developed Rett-like symptoms (20) that progressed rapidly (Fig. 2b), surviving to a median age of 9 weeks (Fig. 2c). P322L males were also underweight compared to littermate controls (Supp. Fig. 3a). These mice were subjected to a program of behavioral tests starting at 6 weeks old when the mutants were only mildly symptomatic. They showed significantly decreased anxiety-like behaviors in the elevated plus maze and open field test (Supp. Fig. 3b,c) although their movement was not significantly impaired (data not shown and Supp. Fig. 3d). The wire suspension test and accelerating rotarod (Supp. Fig 3e,f) exposed severe deficits in motor coordination and learning. Thus P322L males exhibit a severe phenotype mimicking that of *Mecp2*-null mice.

**Figure 2.**
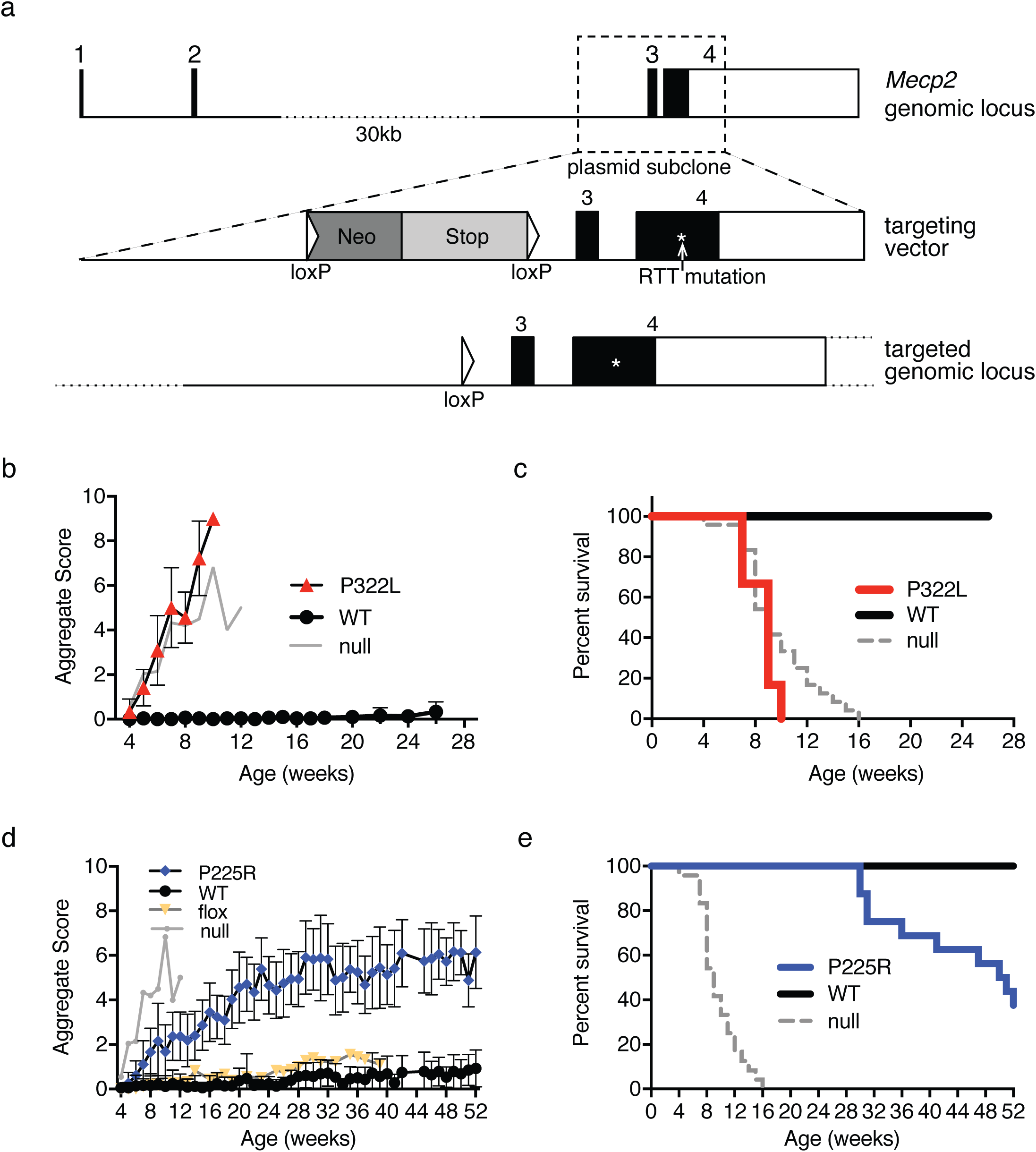
P322L and P225R mutant mice show RTT-like phenotypes and a reduced lifespan. (**a**) The mouse *Mecp2* genomic locus, the targeting vector used to make P225R and P322L knockin alleles in mouse ES cells, and the mutated genomic locus after Cre-mediated removal of the selection cassette. (**b**) P322L hemizygous male mice show a rapid onset of RTT-like phenotypes. Mean aggregate phenotypic score +/-SD plotted for each genotype, P322L n=12 at 4 weeks, WT littermates n=15. Representative *Mecp2*-null data is shown for comparison (n=12 at 4 weeks). (**c**) Kaplan-Meier survival plot for P322L hemizygous males and WT littermates shown in (b). (**d**) P225R hemizygous male mice also display RTT-like phenotypes. Mean aggregate phenotypic score +/-SD plotted for each genotype, P225R n=19 at 4 weeks, WT littermates n=18. Representative *Mecp2*-null and Mecp2-flox data are shown for comparison (flox n=9). (**e**) Kaplan-Meier survival plot for P225R hemizygous males and WT littermates shown in (d).

**Table 1.**
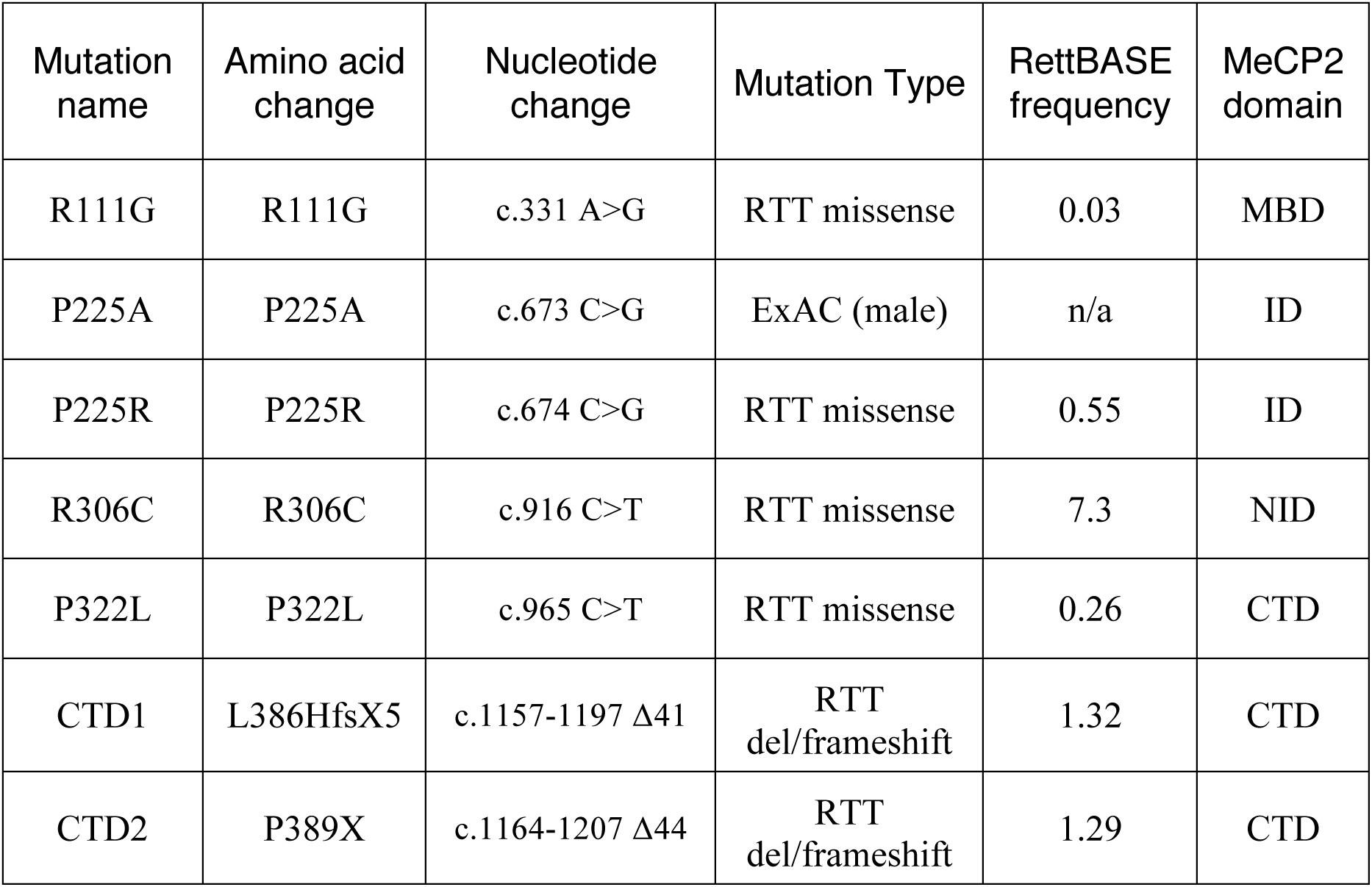
*MECP2* mutations in this study. Amino acid and nucleotide locations are numbered according to the human e2 isoform. RettBASE frequency is expressed as a percentage of all Rett Syndrome cases with an *MECP2* mutation. MBD, methyl-CpG binding domain, ID, intervening domain (between the MBD and NID), NID, NCoR interacting domain, CTD, C-terminal domain.

| Mutation name | Amino acid change | Nucleotide change | Mutation Type | RettBASE frequency | MeCP2 domain |
| --- | --- | --- | --- | --- | --- |
| R111G | R111G | c.331 A>G | RTT missense | 0.03 | MBD |
| P225A | P225A | c.673 C>G | ExAC (male) | n/a | ID |
| P225R | P225R | c.674 C>G | RTT missense | 0.55 | ID |
| R306C | R306C | c.916 C>T | RTT missense | 7.3 | NID |
| P322L | P322L | c.965 C>T | RTT missense | 0.26 | CTD |
| CTD1 | L386HfsX5 | c.1157-1197 Δ41 | RTT del/frameshift | 1.32 | CTD |
| CTD2 | P389X | c.1164-1207 Δ44 | RTT del/frameshift | 1.29 | CTD |

As with P322L mice, P225R males developed Rett-like symptoms at ~6 weeks, but progression of symptoms was more gradual (Fig. 2d). Median survival was 50 weeks (Fig. 2e). P225R males were underweight compared to wild-type littermates (Supp. Fig. 4a). Despite their extended survival time (the longest of any Rett mutation modeled in mice so far), P225R mice showed a similar range of deficits to P322L in behavioral tests. P225R males at 11 weeks of age appeared less anxious than controls in the elevated plus maze and open field test (Supp. Fig. 4b & c). They traveled a shorter distance in the open field (Supp. Fig. 4d) and also demonstrated reduced motor coordination in the wire suspension test (Supp. Fig. 4e), though mutants did not perform less well than controls on the accelerating rotarod (Supp. Fig 4f). We conclude that both P322L and P225R male mice show phenotypes that overlap with those of other mouse models carrying *Mecp2* mutations found in Rett syndrome (21–24).

### P225R and P322L mutations differentially affect MeCP2 levels

To investigate the molecular basis of these RTT-like phenotypes, we first measured MeCP2 abundance. Strikingly, the amount of P322L protein in whole mouse brain was severely reduced (3% of wild-type littermates; Fig. 3a,b). Thus, despite WT amounts of *Mecp2^P322L^* mRNA (Supp. Fig. 5a,b), the level of P322L protein in brain is comparable to that in *Mecp2^Stop/y^* mice, which also show a severe phenotype(22). In P225R mouse brain, MeCP2 protein was at ~22% of the wild-type level (Fig. 3c, d), which is significantly more than in P322L brain. The amount of *Mecp2^P225R^* mRNA was not affected (Supp. Fig 5b).

**Figure 3.**
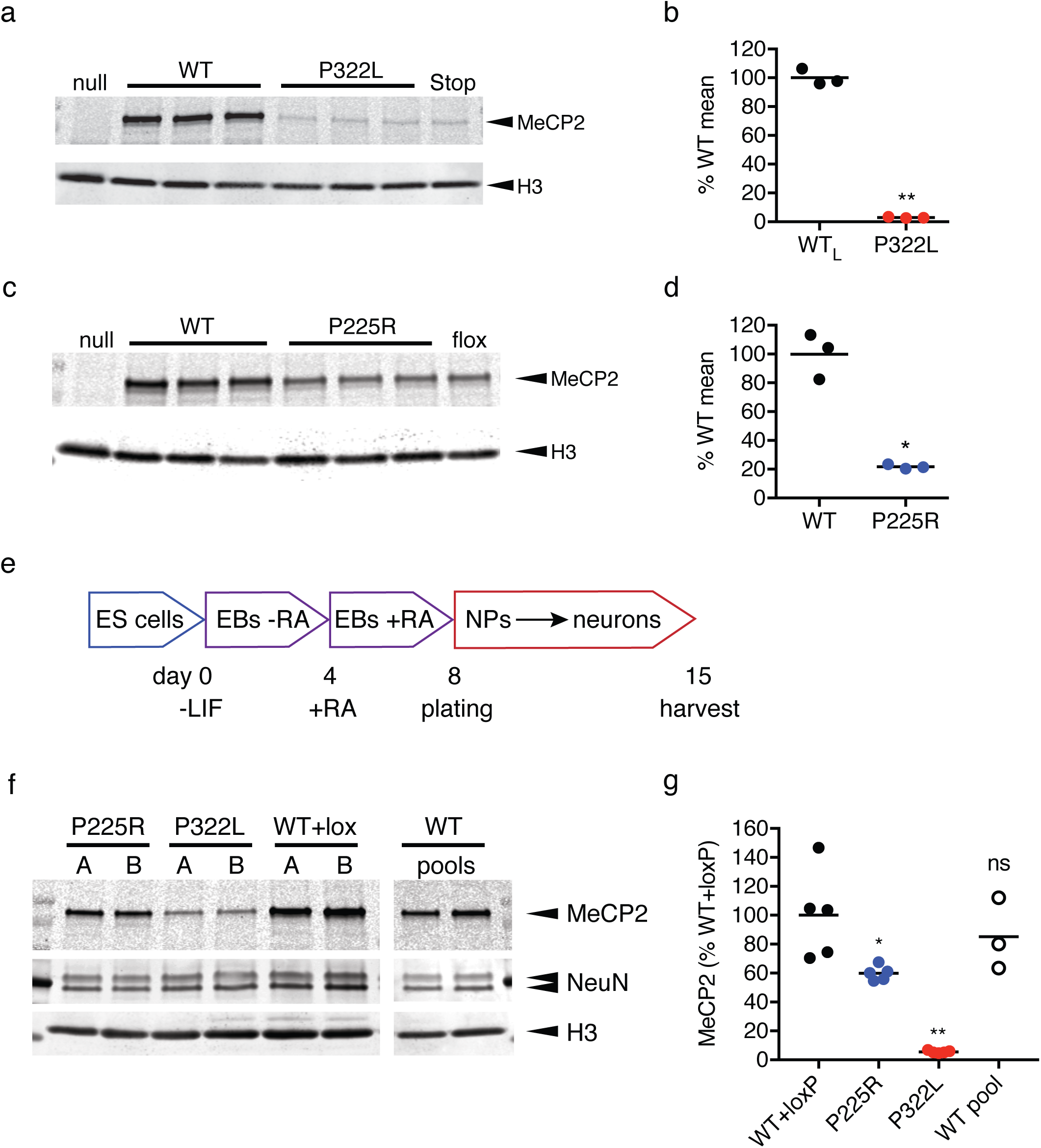
P322L and P225R mutations result in decreased MeCP2 levels in mouse brain and *in vitro* differentiated neurons. (**a**) Western blot of whole brain lysates from 6-week-old P322L and WT littermate males. *Mecp2*-null and *Mecp2*-stop samples are shown for comparison. (**b**) Quantification of (a): the level of MeCP2 (MeCP2 signal/H3 signal) as a percentage of mean WT is shown as mean (black line) and individual biological replicates (n=3 animals). P=0.0010 (**), two-tailed t-test with Welch’s correction for unequal variances. (**c**) Western blot of whole brain lysates from 9-week-old P225R and WT littermate males. *Mecp2*-null and *Mecp2*-lox samples are shown for comparison. (**d**) Quantification of data shown in (c): the level of MeCP2 (MeCP2 signal/H3 signal) as a percentage of mean WT is shown as mean (black line) and individual biological replicates (n=3 animals). P=0.0128 (*), two-tailed unpaired t-test with Welch’s correction for unequal variances. (**e**) Neuronal differentiation protocol used to produce MeCP2-mutant neurons from mouse ES cells. EB: embryoid body, RA: retinoic acid, NP: neuronal precursor (**f**) Representative western blot showing levels of MeCP2 protein present in neurons on day 15 of the differentiation scheme. Lanes A and B for each genotype are derived from independently targeted ES cell clones and WT pools are independent differentiations of the parental ES cell line. Histone H3 was used as a loading control for number of cells and NeuN as a control for equal neuronal differentiation. (**g**) Quantification of MeCP2 levels in *in vitro* differentiated neurons. Two independent clones were differentiated 2 or 3 times for each genotype. MeCP2 level (MeCP2 signal/H3 signal) is normalized to the mean WT+loxP value. Data shown as mean (black line) and individual values. n=5 (WT+loxP, P225R, P322L), n=3 (WT pool) independent differentiations. Comparison to WT+loxP (two-tailed unpaired t-test with Welch’s correction for unequal variances): P225R P=0.0416 (*), P322L P=0.0023 (**), WT (pool) P=0.4868 (ns).

Reduced MeCP2 protein abundance was also observed in cultured neurons derived from the mutant mouse ES cells (Fig. 3e-g). Both P322L and P225R ES cells differentiated equally well into neurons, as shown by NeuN expression (Fig. 3f) and displayed equal amounts of *Mecp2* mRNA (Supp. Fig. 5c). However, protein levels in both mutants were significantly lower than WT and WT+lox controls (Fig. 3f,g). Specifically, P225R showed a moderate reduction to around 60% whereas P322L retained only 7% of the wild-type MeCP2 level.

### P225R and P322L impair NCoR recruitment to chromatin

The severe reduction in P322L MeCP2 explains the severe phenotype, but the P225R mutant is only partially depleted. Notably, the level of P225R MeCP2 in whole brain is similar to that of the hypomorphic floxed *Mecp2* allele (6, 25–27) (Fig. 3c), which shows much milder phenotypic progression than P225R (25–27) (Fig 2d, yellow symbols). To test the possibility that the residual mutant protein is functionally impaired, we looked for other defects that might account for the RTT-like phenotype. While many of the effects of MeCP2 mutations only become apparent when studied in the context of the whole organism, some properties, such as recruitment of the NCoR/SMRT corepressor complex to chromatin, can be tested using a simple transfection assay. P225R and P322L point mutations were introduced into a wild-type mouse *Mecp2* cDNA fused to the C-terminus of EGFP. Interestingly, the ExAC database identifies a hemizygous missense population variant also at P225 (P225A) that presumably does not cause RTT. We included this variant in our analysis and used Rett missense mutations R306C and R111G as negative controls for NCoR and chromatin binding respectively (Table 1). When wild-type EGFP-MeCP2 was expressed in mouse NIH-3T3 cells, green fluorescence co-localized with ‘DAPI bright spots’, which are heterochromatic foci containing high concentrations of methylated DNA (28). A mutation that disrupts the MBD, R111G, abolished binding of MeCP2 to methylated DNA and co-localisation was lost. Both P225R and P322L (in addition to controls P225A and R306C) retained the ability to bind to heterochromatic foci, indicating that MBD function is not affected in these mutants (Fig. 4a).

**Figure 4.**
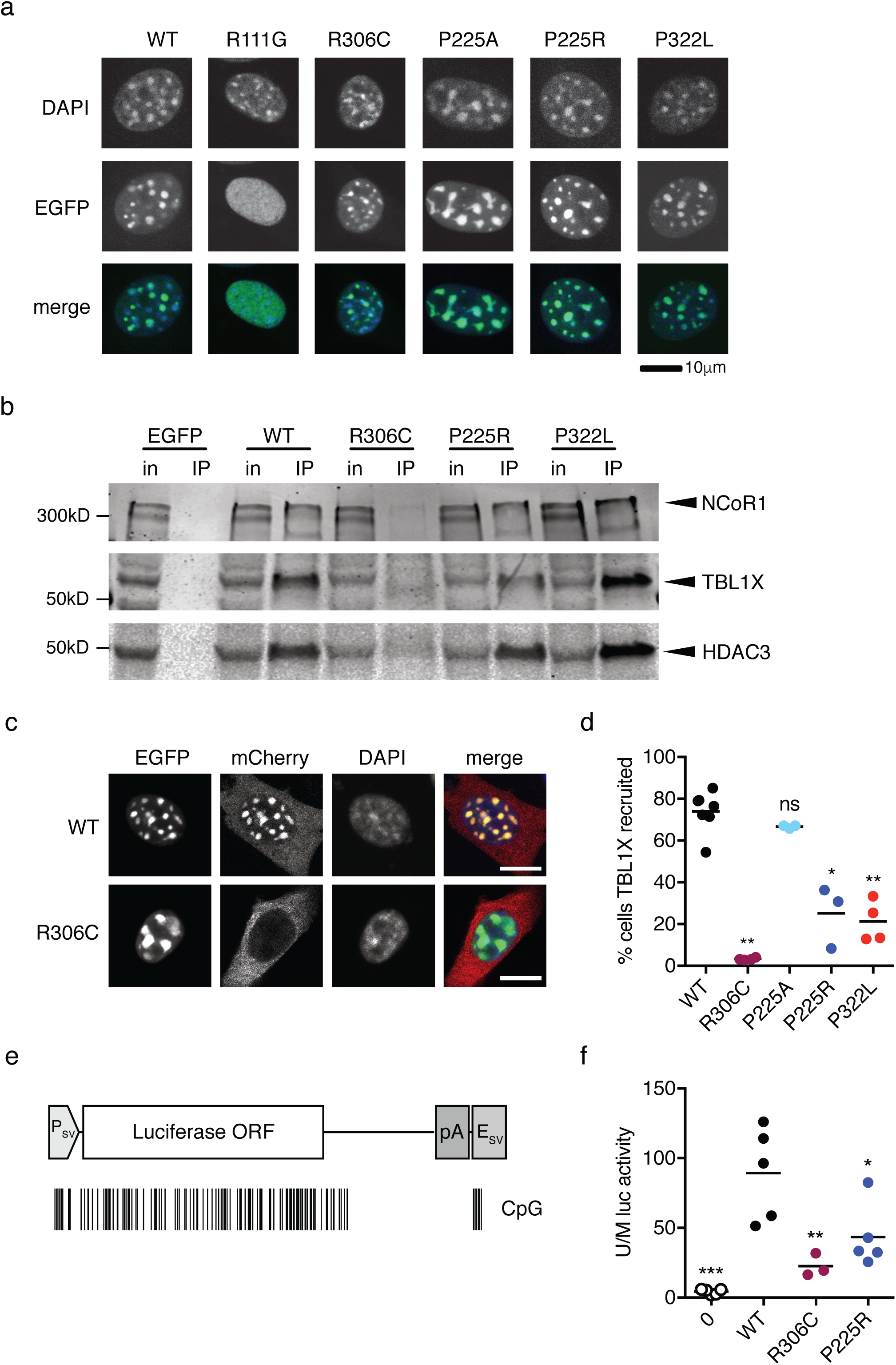
P225R and P322L show impaired NCoR/SMRT recruitment to methylated DNA. (**a**) WT and mutant EGFP-MeCP2 fusions expressed in mouse NIH-3T3 cells. Scale bar 10μm (**b**) Immunoprecipitation of EGFP-MeCP2 from transfected HEK 293 cells with GFP-Trap beads. NCoR/SMRT components NCoR1, TBL1X and HDAC3 detected in western blots of input and immunoprecipitated samples. (**c**) TBL1X recruitment assay. Cotransfection of EGFP-MeCP2 and TBL1X-mCherry expression constructs into NIH-3T3 cells. WT EGFP-MeCP2, but not R306C, recruits TBL1X-mCherry to heterochromatic foci. Scale bar 10μm. (**d**) Quantification of TBL1X recruitment efficiency of MeCP2 mutants. Percent of doubly transfected nuclei with TBL1X-mCherry/EGFP-MeCP2 spots is shown as mean (black line) and individual values. n=7 (WT), 4 (R306C, P322L), 3 (P225A, P225R) independent transfections. MeCP2 mutants were compared to WT using a two-tailed Mann-Whitney test. R306C P=0.0061 (**), P225A P=0.1167 (ns), P225R P=0.0167 (*), P322L P=0.0061 (**). (**e**) Diagram of firefly luciferase expression construct pGL2-Control consisting of an SV40 promoter, *Luc* coding sequence and SV40 polyadenylation signal and enhancer. The positions of CpG dinucleotides methylated by SssI.methylase are shown as black lines. (**f**) Methylation-dependent transcriptional repression assay. Immortalized *Mecp2^−/y^, Mbd2^−/−^* mouse tail fibroblasts transfected with SssI-methylated or unmethylated pGL2-Control plasmid and an MeCP2 expression construct. Methylation-dependent repression expressed as the ratio of unmethylated/methylated (U/M) luciferase activity. For each genotype the mean (black line) and individual values for 3 (R306C) or 5 (no MeCP2, WT, P225R) independent transfection experiments are shown. Genotypes were compared to WT using a two-tailed unpaired t-test with Welch’s correction for unequal variances. No MeCP2 P=0.0045 (***), R306C P=0.0087 (**), P225R P=0.0375 (*).

MeCP2 interacts with the corepressor complex NCoR/SMRT via subunits TBL1X and TBL1XR1. This interaction is disrupted by mutations in the NCoR/SMRT Interaction Domain, including the common Rett mutation R306C. EGFP-tagged MeCP2 was expressed in HEK 293 cells and immunoprecipitated from 150mM salt extracts using GFP-Trap beads (Supp. Fig. 6a). NCoR/SMRT components NCoR1, TBL1X and HDAC3 co-immunoprecipitated with WT MeCP2 but not with the R306C mutant or EGFP alone. Both P225R and P322L mutants were able to immunoprecipitate the same NCoR/SMRT components as the wild-type protein (Fig. 4b), showing that these mutations do not impair the interaction of MeCP2 with the co-repressor complex. The co-repressor mSin3a, which interacts relatively weakly with MeCP2 (29), was immunoprecipitated by all mutant MeCP2 proteins, including R306C (Supp. Fig. 6b).

Although P225R and P322L can still interact with NCoR and methylated DNA separately, we wished to know if they could recruit the NCoR subunit TBL1X to heterochromatin, which requires simultaneous binding to both macromolecules. To test this, constructs expressing both EGFP-tagged MeCP2 and mCherry-tagged TBL1X were transfected into mouse NIH-3T3 fibroblasts (Supp. Fig. 7a-d). It has been shown previously (30) that TBL1X-mCherry, which lacks a nuclear localisation signal, is normally excluded from the nucleus (Supp. Fig. 7b), but in the presence of WT EGFP-MeCP2 it is recruited to the methylation-rich DAPI spots. As expected, the NID mutant R306C itself localizes to the DAPI spots, but is unable to recruit TBL1X (Fig. 4c). When P225R and P322L RTT mutants were tested in this assay both showed impaired recruitment of TBL1X-mCherry to DAPI spots. On the other hand, the population variant P225A performed as well as WT MeCP2 in this assay (Fig. 4d). Thus the ability of both P225R and P322L to recruit NCoR/SMRT to methylated sites in the genome is impaired. Though performed in non-neuronal cells, this test appears to measure a property of MeCP2 that is functionally relevant, as Rett mutations register as defective. Strikingly, substitution of R at position 225 causes Rett syndrome and prevents recruitment, whereas substitution of A at the same site does not cause Rett syndrome and recruits as efficiently as wild-type.

Failure to efficiently recruit the cognate corepressor to methylated sites in the genome suggested that P225R and P322L might also be functionally defective in DNA methylation-dependent transcriptional repression. To test this, untagged MeCP2 was expressed in mouse tail fibroblasts that were compromised in their ability to repress methylated reporter genes due to deletion of the genes for *Mecp2* and *Mbd2* as previously described (6). The co-transfected reporter construct encoding firefly luciferase was either unmethylated or methylated at every CpG (Fig. 4e) and repression was expressed as the ratio of luciferase activities expressed by unmethylated versus methylated constructs. Equivalent expression levels of WT, R306C and P225R MeCP2 constructs were confirmed by western blotting (Supp. Fig. 7e,f). Levels of P322L, however, were consistently low (~30% of WT MeCP2; data not shown), suggesting that lack of an EGFP tag rendered the protein susceptible to degradation, as seen in neurons. As the amount of MeCP2 directly correlates with the strength of repression in this assay, we excluded the P322L mutant from the transcriptional repression experiment. WT MeCP2 repressed the methylated luciferase efficiently, whereas repression by P225R was reduced by about half. The ability of R306C to repress was about 25% that of WT MeCP2 in this assay (fig. 4f). These findings correspond well with the behavior of the mutants in the TBL1X recruitment assay, where R306C was more severely affected than P225R (and P322L), supporting the view that failure to recruit the NCoR corepressor underlies DNA methylation-dependent transcriptional repression by MeCP2. It is possible that the residual repression seen with R306C is due to recruitment of another co-repressor, such as mSin3a, or to direct interference of dense methyl-CpG bound protein with the transcriptional machinery.

### C-terminal deletions do not affect MeCP2 function or recruitment

Deletion-frameshift mutations in the C-terminal domain of *MECP2*, generally between nucleotides c.1100 and c.1200 (human e2 isoform numbering), are responsible for about 10% of all Rett syndrome cases. We chose to model two of the most common confirmed Rett-causing CTDs in mice: c.1157-1197 Δ41 (CTD1) and c.1164-1207 Δ44 (CTD2; see Table 1). Native mouse MeCP2 contains a two amino acid deletion relative to the human protein (Fig. 5a) that coincides with the deletion hotspot in Rett patients. We modeled these Rett mutations in the mouse genome by restoring the two human-specific amino acids and by adding the patient missense tail (CTD1:HQPPX, CTD2:X) (Fig. 5a). We first tested CTD1 and CTD2 for methylated DNA-binding and NCoR/SMRT binding using transfection assays. Both were able to bind to methylated heterochromatic foci in transfected NIH-3T3 cells (Fig. 5b) and to coimmunoprecipitate NCoR/SMRT components NCoR1, TBL1X and HDAC3 (Fig. 5c) and mSin3a (Supp. Fig. 6b). Moreover, unlike P225R and P322L, both CTD1 and CTD2 were able to recruit TBL1X-mCherry to methylated heterochromatic foci as efficiently as WT MeCP2 (Fig. 5d). Thus, neither of the truncated proteins showed any deficiencies in MBD or NID function in these assays.

**Figure 5.**
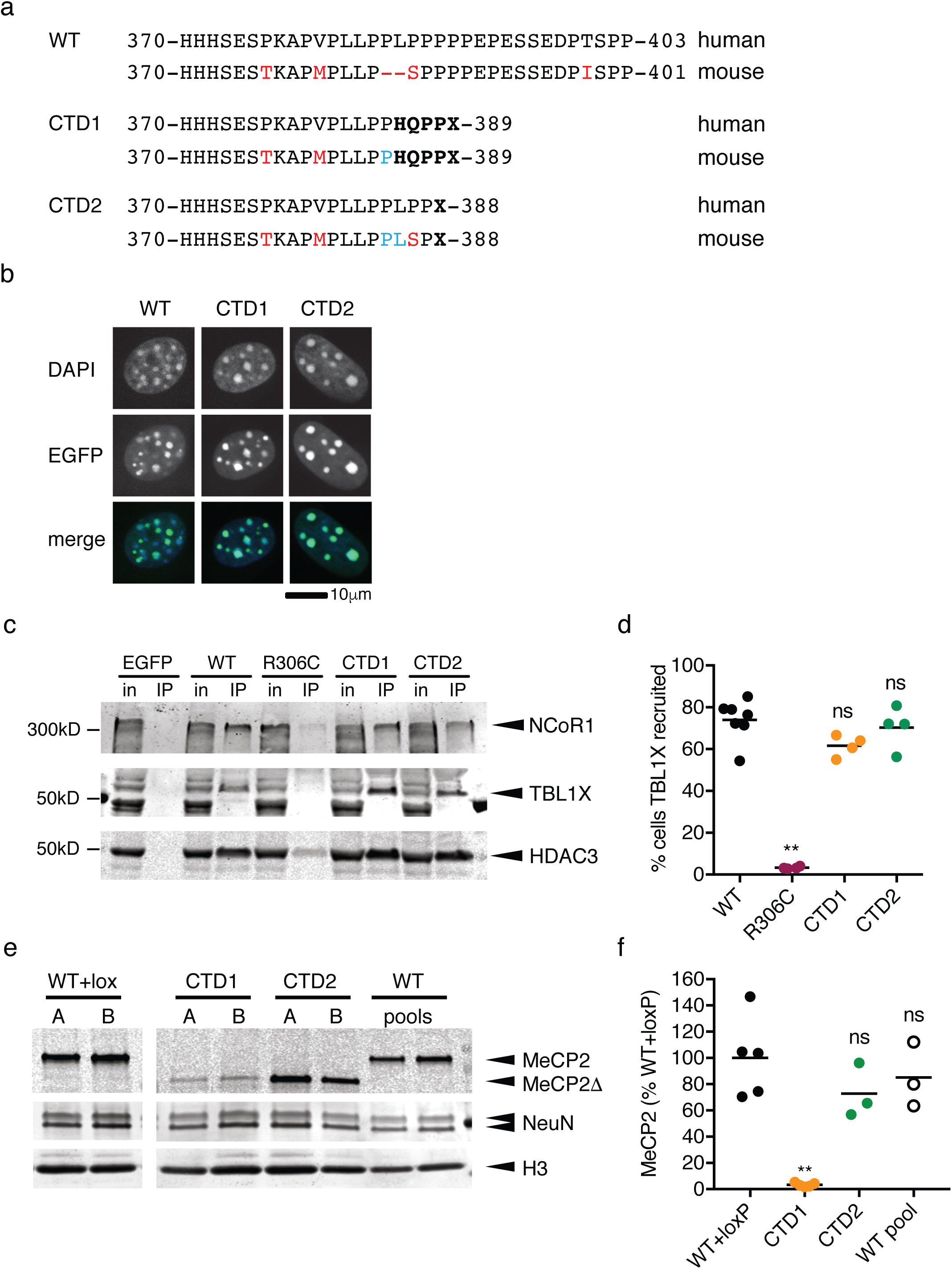
C-terminal truncation mutants CTD1 and CTD2 show normal NCoR/SMRT recruitment but differ in neuronal protein levels. (**a**) Comparison of the human and mouse MeCP2 CTD regions for wild-type (WT) and the patient mutations CTD1 and CTD2. Amino acid changes and absent amino acids in the mouse protein are shown in red, amino acids added to mouse CTD1 and 2 proteins in blue, and missense/nonsense changes arising due to deletion/frameshift in bold type. (**b**) WT, CTD1 and CTD2 EGFP-MeCP2 fusions expressed in mouse NIH-3T3 cells. Scale bar 10μm (**c**) Immunoprecipitation of EGFP-MeCP2 from transfected HEK 293 cells with GFP-Trap beads. NCoR/SMRT components NCoR1, TBL1X and HDAC3 detected in western blots of input and immunoprecipitated samples. (**d**) Quantification of TBL1X recruitment efficiency of MeCP2 mutants. Percent of doubly transfected nuclei with TBL1X-mCherry/EGFP-MeCP2 spots shown as mean (black line) and individual values. WT n=7, R306C, CTD1, CTD2 n=4 independent transfections. Genotypes were compared to WT using a two-tailed Mann-Whitney test. R306C P=0.0061 (**), CTD1 P=0.0727 (ns), CTD2 P=0.6485 (ns). (**e**) Representative western blot showing levels of MeCP2 protein present in neurons on day 15 of differentiation (7 days after plating). Lanes A and B for each genotype are derived from independently targeted ES cell clones and WT pools are independent differentiations of the parental ES cell line. An antibody against the N-terminus of MeCP2 detects both full-length and truncated (Δ) protein. (**f**) Quantification of MeCP2 levels in *in vitro* differentiated neurons. Two independent clones were differentiated 2 or 3 times for each genotype. MeCP2 level (MeCP2 signal/H3 signal) is normalized to the mean WT+loxP value. Data shown is mean (black line) and 5 (WT+loxP, CTD1) or 3 (CTD2, WT pools) independent differentiations. Comparison to WT+loxP (two-tailed unpaired t-test with Welch’s correction for unequal variances): CTD1 P=0.0021 (***), CTD2 P=0.1874 (ns), WT (pool) P=0.4868 (ns).

### CTD1 and CTD2 mutant phenotypes reflect MeCP2 protein level

The absence of overt defects in these assays raised the possibility that CTD1 and CTD2 disrupt other functional domains. To explore this possibility, we used CTD1 and CTD2 targeting vectors that retained all sequences downstream of the deletions to create *Mecp2^CTD1/y^* and *Mecp2^CTD2/y^* ES cell lines (Supp. Fig 8a,b). After Cre-mediated deletion of the floxed Neo-Stop selection cassette, the mutated *Mecp2* alleles differed from WT only at the mutation site and a loxP site upstream of exon 3. When these lines were differentiated into neurons *in vitro,* both genotypes produced truncated MeCP2 protein as expected, but surprisingly the amount of protein differed greatly. CTD1 retained only 3% the WT level, whereas the amount of CTD2 MeCP2 was not significantly different from WT controls (Fig. 5e,f). At the transcript level, CTD1 had only 34% WT mRNA whereas CTD2, like P225R and P322L, expressed WT-levels of mRNA (Supp. Fig. 8c).

The extremely low level of MeCP2 protein in CTD1 neurons predicted that mice would show a severe phenotype, similar to P322L and *Mecp2*-null mice. To test this, we produced CTD1 mice using CRISPR/Cas9 cutting and repair with an oligonucleotide template in mouse zygotes. After confirming the presence of the mutation by sequencing (data not shown) and Southern blot (Supp. Fig. 9a,b), analysis of CTD1 founder males showed that the amount of MeCP2 in whole brain was reduced to about 10% of that in WT controls (Fig 6a, b) and the mRNA to 45% of WT (Supp. Fig. 9e). A cohort of CTD1 male mice was scored for Rett-like phenotypes, which progressed rapidly (Fig. 6c), giving a median survival of 20 weeks (Fig. 6d). The weights of CTD1 mice were not significantly different to those of WT littermates (Supp. Fig. 9c). This cohort was still on a mixed CBA/C57BL6 genetic background whereas small size is usually seen in *Mecp2*-mutant males on a more inbred C57BL6 background.

**Figure 6.**
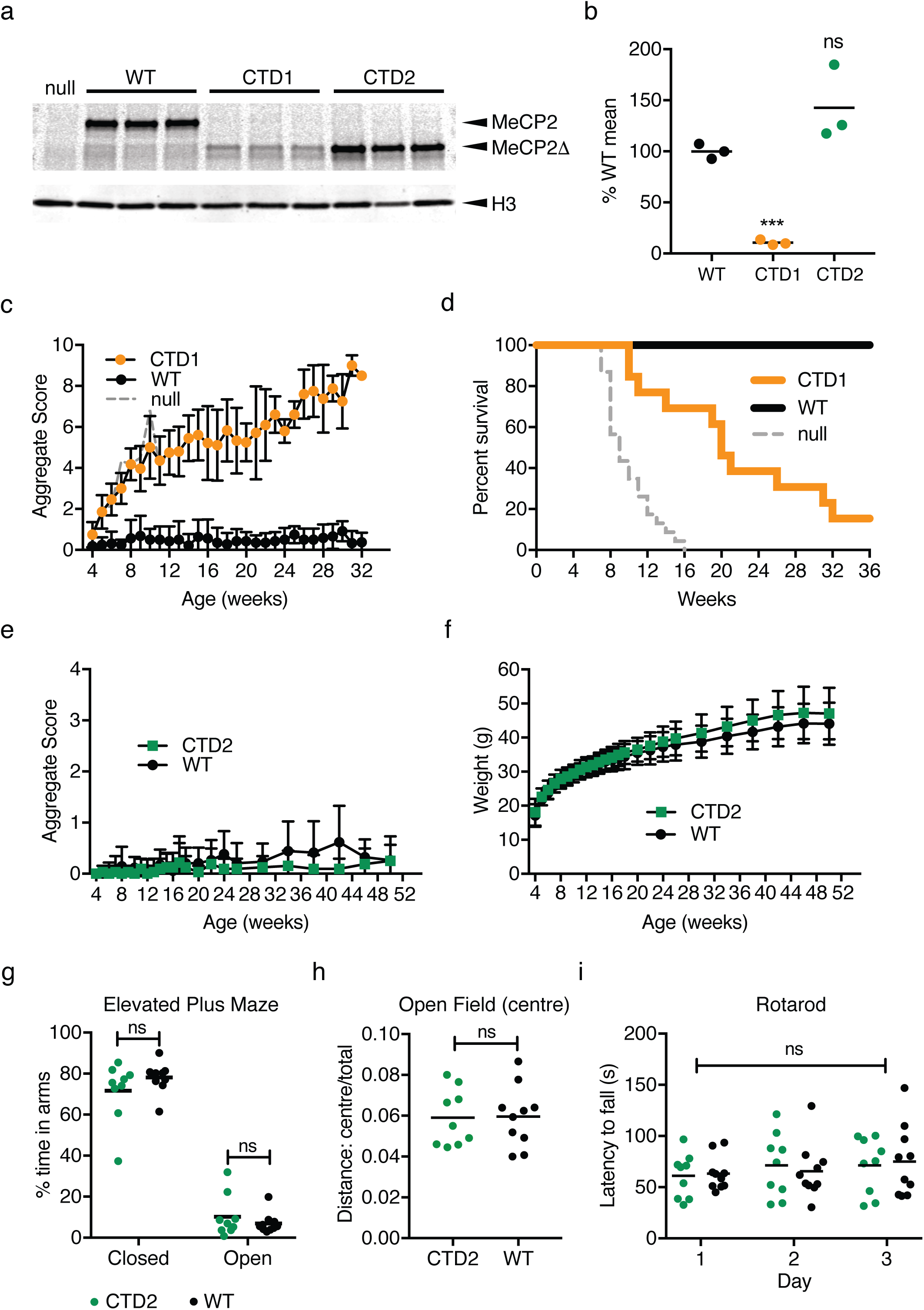
CTD1 and CTD2 mice show contrasting phenotypes due to different levels of MeCP2 in brain. (**a**) Western blot of whole brain lysates from 6-week-old CTD1, CTD2 and WT males. A *Mecp2*-null sample is shown for comparison. CTD1 and CTD2 mice express a truncated protein, MeCP2Δ. (**b**) Quantification of (a): the level of MeCP2 (MeCP2 signal/H3 signal) as a percentage of mean WT shown as mean (black line) and individual values for each genotype (n=3 animals). Comparison to WT (two-tailed unpaired t-test with Welch’s correction for unequal variances): CTD1 P=0.0008 (***), CTD2 P=0.1773 (ns). (**c**) CTD1 hemizygous males display RTT-like phenotypes. Mean aggregate phenotypic score +/-SD plotted for mutants and WT male littermates, CTD1 n=11, WT n=7 (at 4 weeks). (**d**) Kaplan-Meier survival plot for CTD1 hemizygous males and WT littermates shown in (c). (**e**) Phenotypic scoring of CTD2 males and WT littermates. Mean aggregate phenotypic score +/-SD plotted for mutants and WT male littermates, CTD2 n=16 WT n=17. (**f**) Mean weight +/-SD for animals shown in (e). (**g**) Elevated plus maze. Percentage of time spent in the closed and open arms is shown as mean (black line) and individual values. Percentage time in open and closed arms for each genotype was compared using two-tailed unpaired t-tests: closed arms P=0.2238 (ns), open arms P=0.3713 (ns). (**h**) The ratio of distance travelled in the central zone of the open field arena to total distance travelled shown as mean (black line) and individual values. Two-tailed unpaired t-test test P=0.9394 (ns). (**i**) Accelerating rotarod test. The latency to fall is shown as mean (black line) and individual data points on each day. Two-way ANOVA (repeated measures) [genotype effect, F(1, 17) = 9.451×10^−6^, P=0.9976 (ns)]. (g)-(i) CTD2 n=9, WT n=10.

The instability of CTD1 made it impossible to deduce whether the C-terminal truncation mutation gave rise to a novel functional defect in MeCP2 as at 10% of WT littermate controls the level of protein was similar to the level of full-length protein in *Mecp2^Stop/y^* mice, which resemble *Mecp2*-null mice (22). On the other hand, the fully stable CTD2 mutant protein offered the chance to test the importance of this C-terminal region. CTD2 mice were therefore produced using standard injection of ES cells into mouse blastocysts and verified by Southern blots (Supp. Fig. 9d) and sequencing (data not shown). As in cultured neurons, CTD2 MeCP2 protein and mRNA levels in whole brain were not significantly different from wild-type littermate controls (Fig. 6a,b, Supp. Fig 9e). Strikingly, CTD2 male hemizygous mice and WT littermate controls were phenotypically indistinguishable over the course of one year (Figure 6e,f). Survival was 100% and CTD2 mice behaved as WT in the elevated plus maze (Fig. 6g), open field test (Fig. 6h, Supp. Fig. 9f) and accelerating rotarod (Fig. 6i). In the wire suspension test CTD2 mice performed significantly better than WT littermates (Supp. Fig. 9g). We conclude that the CTD2 C-terminal truncation of MeCP2 has no overt phenotypic consequences in this mouse model and therefore does not identify a previously unknown functional domain.

### ‘Humanisation’ of the CTD2 allele reduces protein levels in neurons

All Rett mutations previously studied in mice have replicated the disease seen in patients, including the relative severity of symptoms (5, 6, 23, 24, 31–33). Why should CTD2 be exceptional? A potential explanation was that subtle differences in amino acid sequence between human and mouse MeCP2 proteins were involved. Despite 95% identity of MeCP2 protein between the two species, several amino acid differences persisted near the truncated region (Fig. 7a). To test whether these differences altered stability of the mutant, we created a “humanised” CTD2 allele (CTD2hu) in mouse ES cells by making three single nucleotide alterations: T376P, M380V and S387P (Fig. 7a). These changes ensure that the amino acid sequence of the C-terminus of MeCP2 in patient and mouse alleles is identical from E298, which is N-terminal to the NID, to the stop codon. CTD2hu ES cell clones were produced (Supp. Fig. 10a,b) and differentiated into neurons for comparison with CTD1, CTD2 and WT+lox controls. Western blots showed a dramatic destabilisation of CTD2hu, which was now indistinguishable from CTD1 (7.4 and 5.0% WT+lox respectively Fig. 7b,c). The original CTD2 MeCP2 protein remained essentially identical to WT+lox controls (93%). CTD2hu neurons also contained less mRNA than WT+lox and CTD2, again resembling CTD1 (Supp. Fig. 10c). Thus, altering the sequence context of the non-pathogenic CTD2 mouse allele to make it resemble the patient mutation more closely resulted in a drastic reduction in the amount of MeCP2 in neurons.

**Figure 7.**
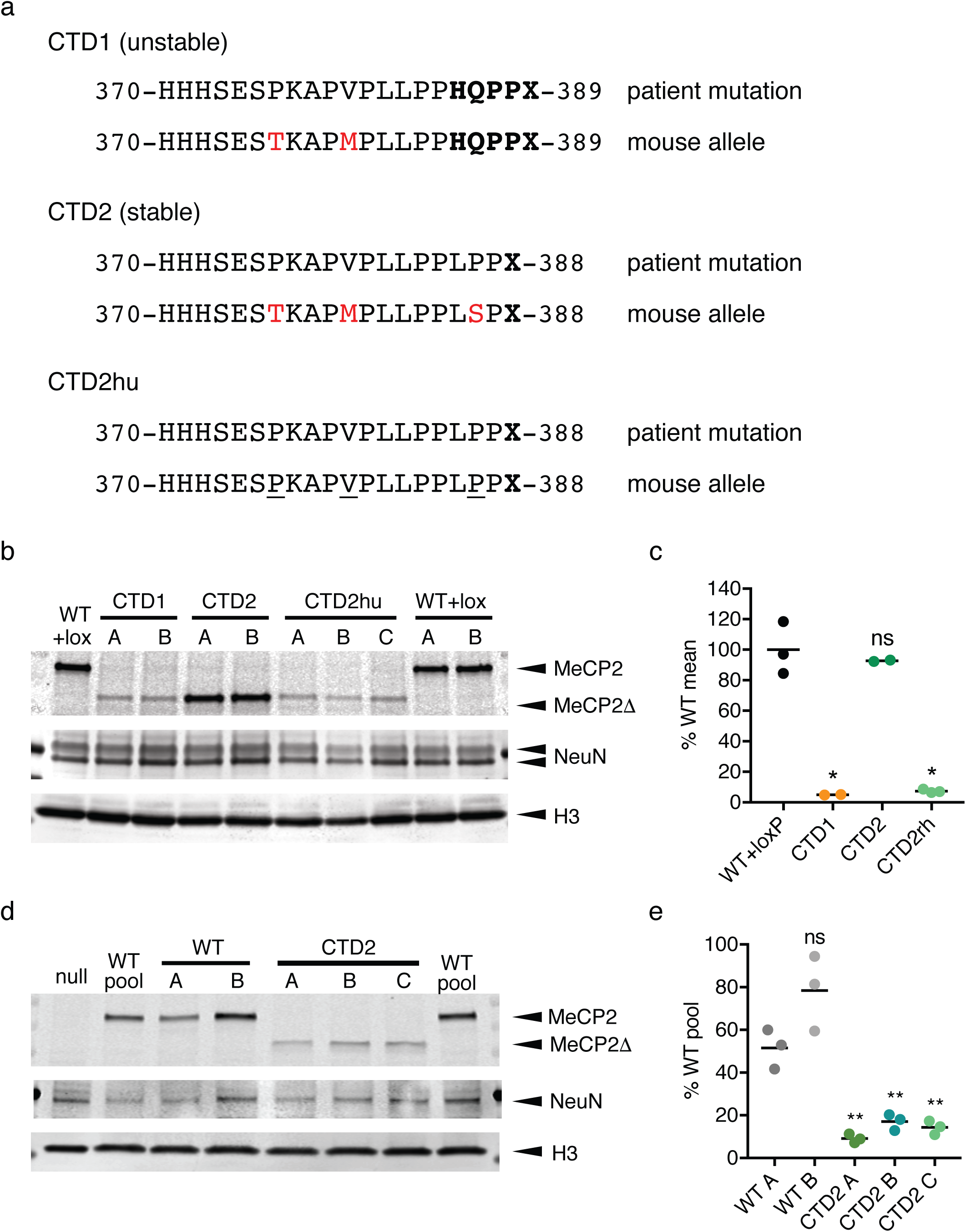
A humanized CTD2 allele expresses low levels of truncated MeCP2. (**a**) The amino acid sequences of the extreme C-termini of CTD1, CTD2 and CTD2hu mouse alleles and their corresponding human RTT alleles. Missense/nonsense changes caused by the deletion-frameshift are in bold type. Residues where the mouse allele differs from human are shown in red. Changes between CTD2 and CTD2hu are underlined. (**b**) Western blot showing levels of MeCP2 protein present in neurons on day 15 of differentiation (7 days after plating). Lanes A,B (and C) for each genotype are derived from independent mouse ES cell clones. An antibody against the N-terminus of MeCP2 detects both full-length and truncated (Δ) protein. Quantification of MeCP2 levels in *in vitro* differentiated neurons. Two or three independent clones were differentiated for each genotype. MeCP2 level (MeCP2 signal/H3 signal) is normalized to the mean WT+loxP value. Data shown is mean (black line) and individual values for 3 (WT+loxP, CTD2hu) or 2 (CTD1, CTD2) independent differentiations. Comparison to WT+loxP (two-tailed unpaired t-test with Welch’s correction for unequal variances): CTD1 P=0.0108 (*), CTD2 P=0.5450 (ns), CTD2hu P=0.0111 (*). Western blot showing levels of MeCP2 protein present in LUHMES-derived human neurons on day 9 after precursor plating. Differentiations of 1 MECP2-null clone, 2 WT pools, 2 unmodified WT clones and 3 CTD2 LUHMES clones are shown. An antibody against the N-terminus of MeCP2 detects both full-length and truncated (Δ) protein. (**e**) Quantification of MeCP2 levels in LUHMES-derived neurons. Three independent differentiations were performed for each of the clones shown in (d). MeCP2 level (MeCP2 signal/H3 signal) is normalized to the WT pool value for each differentiation. Data shown is mean (black line) and individual values for 3 independent differentiations. Comparison to WT A (two-tailed unpaired t-test): WT B P=0.0798 (ns), CTD2 A P=0.0015 (**), CTD2 B P=0.0039 (**), CTD2 C P=0.0027 (**).

To further confirm our hypothesis that CTD2 MeCP2 is unstable in human neurons we created a knock-in *MECP2^CTD2^* allele in LUHMES cells, a human neuronal progenitor cell line derived from human embryonic ventral mesencephalic tissue which can be differentiated *in vitro* to produce a uniform population of mature dopaminergic neurons(34, 35). Knock-in clones containing *MECP2^CTD2^* alleles on the active X chromosome, and thus expressing MeCP2 CTD2 protein, were produced using CRISPR/Cas9 cutting and an oligodeoxynucleotide repair template as previously described (35) (Supp. Fig. 10d,e). After differentiating for 9 days to mature neurons, the level of CTD2 MeCP2 protein was severely reduced compared to the WT parental cells and unmodified WT clones (Fig. 7d,e). MeCP2 protein levels in CTD2 clones ranged from 7-20% of the parental WT cells. These findings in mice, mouse ES cell-derived neurons and human neurons strongly suggest that Rett syndrome resulting from C-terminal deletion mutations is due to severe MeCP2 deficiency rather than the loss of essential domains in the C-terminal region.

## Discussion

Biochemical and cell biological studies of MeCP2 have identified numerous interacting proteins that potentially mediate its function. So far, only two of these are implicated in Rett syndrome, as causal missense mutations disrupt the interactions with methylated DNA or with the NCoR/SMRT corepressor complexes (11). We were intrigued by a number of RTT mutations that leave the MBD and the NID intact, and by the possibility that they might direct us to additional functional domains that had previously escaped detection. To be certain that the mutations are genuinely causative of the disorder, we confined our attention to variants shown to be absent in parental DNA. The importance of this criterion is illustrated by E397K, which has been detected several times in classical RTT (7), but has subsequently been classified as a relatively common and phenotypically benign population variant (7, 8). By studying C-terminal truncations and the missense mutations P225R and P322L, which are recurrent RTT mutations remote from the two known interaction sites, we failed to establish new functional domains of MeCP2. Instead, our results can provide coherent molecular explanations for the deleterious effects of these mutations by considering MeCP2 as a recruiter of histone deacetylase complexes to chromatin (29). Further evidence that it is the conformation and/or stability of MeCP2 that is affected by these mutations rather than functional domains containing P225, P322 or the CTD region comes from a severely truncated mouse knock-in allele, *ΔNIC*, which despite lacking all of these sites shows a very mild phenotype, distinct from that seen in RTT mouse models (14).

It is clear from studies of patient severity (18, 33, 36, 37) and mouse models (5, 6, 24, 31) that mutations in the MBD and the NID, such as T158M and R306C, are somewhat less severe on average than complete absence of the protein. This suggests that these mutant proteins retain some function, which could come from a number of sources. T158M does not completely abolish binding to methylated DNA (24) and some residual binding of R306C to NCoR components can be seen in this study (Fig. 4b, Supp. Fig 6b). All MeCP2 proteins tested could bind to mSin3a (Supp. Fig. 6b). It has been proposed that other domains such as AT-hooks (38, 39) may also contribute low-level function of MeCP2 when the MBD and NID are mutated.

P225R leads to moderate destabilisation of MeCP2, but it also interferes with its ability to recruit the co-repressor subunit TBL1X to chromatin and to impose DNA methylation-dependent transcriptional repression. Although P225 is in an unstructured region of the protein, it has been reported that conformational changes occur on binding to DNA (40). P225R may hinder the ability of MeCP2 to form the required bridge between chromatin and corepressor complex. It is also possible that the same conformational change leads to instability of the protein. The combination of these two partial defects in stability and function can explain the moderate RTT-like phenotype of these mice. Importantly, a rare human population variant at the same amino acid, P225A, does not interfere with co-repressor recruitment. The strong correlation between the ability to recruit co-repressor to methylated DNA and phenotypes in mice and humans strengthens the notion that this is a key role of MeCP2.

The other non-canonical missense mutation, P322L, severely destabilises MeCP2 and this alone can account for the resulting severe phenotype. The C-terminal truncations CTD1 and CTD2hu also severely destabilise the protein in neurons, but do not detectably compromise several aspects of its function in cell transfection-based assays. Emphasising this point, CTD2 mice, which lack 100 amino acids from the C-terminal end of MeCP2, are viable, fertile and phenotypically normal in a series of behavioral assays. We note that this truncation removes an activity-dependent phosphorylation site at S421 (S423 in human MeCP2) that has previously been implicated in neuronal function, including dendritic patterning and spine morphogenesis (41–44). The region has also been implicated in regulation of micro-RNA processing and dendritic growth (45). Although loss of these functional domains evidently does not make a detectable contribution to the major RTT-like phenotypes in mice, we cannot exclude the possibility that subtle phenotypic consequences were not detected by our assays or that they are important for higher functions present in humans but not in mice.

MeCP2 instability has previously been noted as a result of mutations that also disrupt the functional domains of MeCP2, including missense mutations T158A (31), T158M and R133C (24) and nonsense mutations R168X (23) and R255X (32). Our results show that MeCP2 deficiency due to protein destabilisation alone is a major underlying cause of RTT. Although we only tested two CTD mutations, we suggest that severe MeCP2 destabilisation is a mechanism that applies generally to this RTT mutation category. Why mutations outside the key functional domains should confer instability is unknown. With the exception of the MBD and NID domains, whose 3D conformation has been determined (12, 13), most of the protein is thought to be unstructured (40). Surprisingly, the instability of CTD2 in the context of human MeCP2 was suppressed by changing three nearby amino acids to match the mouse MeCP2 sequence. This serendipitously allowed us to determine that the deletion of the MeCP2 CTD beyond amino acid 388, thereby losing ~100 amino acids from the C-terminus, has no deleterious phenotypic consequences. We conclude therefore that this mutation does not identify a novel functional domain, but causes RTT in humans purely due to drastic destabilisation of the protein.

The dramatic difference in the stability of CTD2 conferred by human versus mouse sequences indicates that subtle changes in protein primary structure can have dramatic consequences for MeCP2 abundance. It is also interesting that CTD alleles that result in reduced protein levels also have less mRNA, although the decrease is less severe. Further investigation of the precise cause of CTD instability will be a focus of future research.

Importantly, the finding that CTD mutations do not appear to affect MeCP2 function may offer therapeutic opportunities. A recent study has demonstrated the use of proteasome inhibitors to increase the level of unstable T158M MeCP2 (46). Alternatively, small molecules that bind to the mutant CTD and affect its conformation may improve stability and therefore provide clinical benefit.

## Materials and Methods

### *Mecp2* mutant mouse alleles

RTT mutations P225R, P322L, CTD1, CTD2 and CTD2hu were introduced into a targeting vector (20) (Fig. 1a) containing WT *Mecp2* sequences and a Cre-excisable selection cassette using the QuikChangeII XL Site-Directed Mutagenesis kit (Agilent Technologies). Targeting vectors were linearized at the 3’ end using NotI and electroporated into 129/Ola E14Tg2a mouse ES cells (A gift from A.Smith, University of Edinburgh). Correctly targeted clones were identified by an initial PCR screen:

> forward primer: 5’-TCACCATAACCAGCCTGCTCGC-3’
>
> reverse primer: 5’-ATTCGATGACCTCGAGGATCCG-3’)

followed by Southern blotting of candidate clones and sequencing of the mutation sites (Supp. Fig. 1, 7). A number of clones that had recombined to include the floxed selection cassette but no RTT mutation were also verified for use as controls (WT+lox).

### Differentiation of ES cells into neurons

ES clones were transiently transfected with a pCAGGS-Cre expression plasmid and clones that had deleted the Neo selection cassette were verified by PCR, Southern blotting and sequencing before differentiation into neurons. Two independently targeted clones were used for each genotype, and each clone was differentiated into neurons two or three times.

Differentiation was carried out using a 4-/4+ retinoic acid (RA) procedure as previously described (47, 48). Neural precursor cells were seeded onto 6cm dishes at a density of 1.5×10^5^ cells/cm^2^ and harvested after 7 days by scraping into phosphate-buffered saline (PBS), pelleting the cells and snap freezing.

### Generation of CTD2 LUHMES cell lines

The LUHMES cell line (ATTC CRL-2927) was obtained from ATCC and cultured according to the methods described in Scholz *et al* (49), with some minor alterations. All vessels were coated in poly-L-ornithine (PLO) and fibronectin overnight at 37°C. Proliferating LUHMES cells were seeded at 2×10^6^ cells/T75 flask every 2 days. For differentiation, 2.5×10^6^ cells were seeded in a T75 flask for the first 2 days of the protocol and on day 2 cells were seeded at 6×10^6^cells/10cm dish. During differentiation a half-media change was performed on day 6 and neurons were harvested for protein on day 9.

To introduce a CTD2 knock-in mutation, LUHMES cells were transfected with plasmid pSpCas9(BB)-2A-GFP (pX458) (a gift from Feng Zhang, Addgene plasmid #48138) containing the sgRNA sequence 5’-TCCTCGGAGCTCTCGGGCTC-3’, and single-stranded oligodeoxynucleotide 5’CCATCACCACCACTCAGAGTCCCCAAAGGCCCCCGTGCCACTGCTCCCACCCCTGCCGCCCTGAGCCCCAGGACTTGAGCAGCAGCGTCTGCAAAGAGGAGAAGATGCCCAGAGGAGGCT3’ (Sigma, desalted). Cells were transfected by Nucleofection (Lonza) using a Basic Nucleofector kit for primary neurons (VAPI-1003) and a Nucleofector II device, and FACS-sorted to isolate clones as previously described (35).

LUHMES clones were analysed by sequencing of genomic DNA and cDNA and by Southern blotting (Supp. Fig. 10d,e) to isolate clones that were either homozygous for the CTD2 mutation, or heterozygous and expressing the CTD2 mutation from the active X-chromosome (LUHMES cells are female, XX). Three CTD2 clones (two homozygous and one heterozygous) and two unmodified WT control clones were used for further analysis.

LUHMES clones were differentiated to mature neurons as previously described (49). A null clone (H4) was described previously (35). All clones and the parental cell line underwent three independent differentiations and were harvested on day 9 of differentiation by scraping into phosphate-buffered saline (PBS), pelleting the cells and snap freezing.

### Establishment of *Mecp2* mutant mouse lines

P225R, P322L and CTD2 mouse lines were generated by injection of ES cells into mouse blastocysts using standard methods. Chimeric males were crossed with CMVCre deleter females (50) to remove the selection cassette. This and subsequent generations were bred by crossing *Mecp2-*mutant heterozygote females with C57BL/6J WT males. The genotypes of mutant lines were confirmed by Southern blotting (Supp. Fig. 3a,b, 8d) and sequencing. Routine genotyping was performed by PCR across the loxP “scar” site remaining in the mutant alleles after selection cassette removal (forward primer: p5 5’-TGGTAAAGACCCATGTGACCCAAG-3’, reverse primer: p7 5’-GGCTTGCCACATGACAAGAC-3’, WT 416bp, WT+loxP 558bp).

The CTD1 mouse line was created by pronuclear injection of Cas9 protein (Integrated DNA Technology), synthetic tracrRNA and crRNA (target sequence ACCTGAGCCTGAGAGCTCTG) (Sigma) and an oligonucleotide repair template (GCACCATCATCACCACCATCACTCAGAGTCCACAAAGGCCCCCATGCCACTGCTCCCACCCCACCAGCCCCCCTGAGCCTCAGGACTTGAGCAGCAGCATCTGCAAAGAAGAGAAGATGC) (Sigma) into C57BL6/J:CBA/CaOlaHsd F2 fertilized oocytes as previously described (51). Correctly mutated founder animals were identified by sequencing across the deletion site. One CTD1 hemizygous male founder was able to breed and female heterozygous offspring were checked by sequencing and Southern blot (Supp. Fig. 8a,b) and used for further breeding of the line.

### Phenotypic analysis

Cohorts of hemizygous mutant male mice and WT littermates were weighed and scored weekly for a range of RTT-like phenotypes to give an aggregate score between 0 and 12 as previously described (20, 25). Cohorts of at least 8 mutants and 8 wild-type littermates were scored for each genotype, with higher numbers being used where available to allow for any losses unrelated to the mutation, such as fighting. Scoring was carried out blind to genotype and to previous scores. Animals that scored a maximum score of 2 for tremor, breathing or general condition or which had lost 20% of their body weight had reached the severity limit of the experiment according to the Home Office license and were humanely culled. These animals were counted as having ‘died’ for the purposes of survival data. Animals of any genotype which were culled for reasons not linked to the mutation, such as fighting with cage mates, were removed from survival plots at that point (censored data). P225R, P322L and CTD2 animals had been backcrossed to C57BL6/J for 3 generations and CTD1 animals only once.

Behavioral testing was carried out on cohorts of mice that had been backcrossed to C57BL6/J for 4 generations. Testing took place over a 9-day period in the order: day 1: elevated plus maze (1×15 minute trial), day 2: open field test (1×20 minute trial), day 3: wire suspension test (3 trials of 30 seconds separated by 15 minutes), day 6: accelerating rotarod habituation (5 minutes at 4rpm, days 7-9: accelerating rotarod trials (4 trials per day separated by 1 hour). Tests were performed as previously described(24, 25). Hemizygous male mutant mice and WT littermates were tested at an age appropriate to the development of symptoms for that line: testing was started for P322L at 6 weeks, P225R 11 weeks and CTD2 18 weeks. A cohort size of 10 animals per genotype was chosen as the largest number of animals that could reasonably be tested in a single session. Mice were housed in mixed mutant and wild-type littermate groups. Mice were tested blind to genotype and in order of ID number, giving a random order of mutant and WT animals. Positions on the rotarod were assigned to ensure equal representation of mutant and control animals at each of the five positions.

### RNA preparation and qRT-PCR

Total cellular RNA was prepared from *in vitro* differentiated neurons (7 days after plating, 10^7^ cells) and mouse brain (half brain) using TriReagent (Sigma). cDNA was prepared using a QuantiTect kit (Qiagen) and amplified in a qPCR reaction using SensiMix SYBR and Fluoroscein Master Mix (Bioline) using primers for *Mecp2* (forward: 5’-ACCTTGCCTGAAGGTTGGAC-3’, reverse: 5’-GCAATCAATTCTACTTTAGAGCGAAAA-3’) and control *Cyclophilin A* (forward: 5’-TCGAGCTCTGAGCACTGGAG-3’, reverse: 5’-CATTATGGCGTGTAAAGTCACCA-3’). Samples were run in triplicate and the amount of *Mecp2* cDNA calculated for each biological replicate using the ‘double delta’ method after correction of C_T_ values using a standard curve.

### Protein extracts and western blotting

*In vitro* differentiated ES cell-derived and LUHMES-derived neurons and whole brain samples were prepared for western blotting as previously described (3). Extracts were run on TGX 4-20% gradient gels (BioRad) loading extract equivalent to approximately 5 x 10^5^ nuclei per well. Samples were run on duplicate gels and transferred to nitrocellulose membrane overnight in the cold at 25V. Western blots were processed as described previously (3) using the following antibodies: anti-MeCP2 (N-terminus): mouse monoclonal Men-8 (Sigma), anti-NeuN: rabbit polyclonal ABN78 (Millipore), anti-histone H3: rabbit polyclonal ab1791 (Abcam). Western blots were developed with IR-dye secondary antibodies (IRDye 800CW donkey anti-mouse, IRDye 680LT donkey anti-rabbit, LI-COR Biosciences) and scanned using a LI-COR Odyssey machine. Images were quantified using Image Studio Lite software (LI-COR Biosciences). The ratio of MeCP2:NeuN signals for each lane was used to check for equal neuronal *in vitro* differentiation and clones that did not differentiate well were discarded. MeCP2 levels were normalized to the histone H3 signal for each lane to compare the amount of MeCP2/nucleus between samples.

### EGFP-tagged cDNA constructs and transfection assays

Mouse *Mecp2* cDNA (e2 isoform) was cloned into the vector pEGFP-C1 (Clontech) to create in-frame fusions with an N-terminal EGFP tag. Missense mutations were introduced using the QuikChangeII XL Site-Directed Mutagenesis kit (Agilent Technologies) and truncations by PCR amplifying the cDNA with a modified reverse primer before cloning into the expression vector. A vector expressing mCherry-tagged mouse TBL1X was described previously(30). For localisation of EGFP-MeCP2 and colocalisation of EGFP-MeCP2/TBL1X-mCherry constructs were transfected into NIH-3T3 mouse fibroblasts (93061524, ECACC) growing on glass coverslips using Lipofectamine (Thermo Fisher). Cells were fixed in 4% Paraformaldehyde 48 hours after transfection, stained with 4’,6-Diamidino-2-Phenylindole (DAPI) and mounted in ProLong Diamond mountant (Thermo Fisher). Images were captured using a Leica SP5 confocal microscope with 63x objective. For western blotting to test expression levels of EGFP-MeCP2 WT and mutant proteins cells were transfected with EGFP-MeCP2 and TBL1X-mCherry expression constructs as above and harvested 48 hours after transfection by trypsinisation. Cell pellets were prepared for western blotting as described above for *in vitro* differentiated neurons. MeCP2 was detected using mouse monoclonal Men-8 (Sigma) and anti-γ-tubulin mouse monoclonal GTU-88 (Sigma) was used as a loading control.

For quantification of TBL1X-mCherry colocalisation with EGFP-MeCP2 3-7 independent transfections were performed for each genotype. Transfected cells were selected first for their EGFP signal, without observing the mCherry channel, and then scored for colocalisation of the mCherry signal. In each transfection at least 15 cells on 4 coverslips (total 60 cells per genotype) were counted, blind to the genotype.

### Coimmunoprecipitation assay

EGFP-MeCP2 expression constructs were transfected into HEK 293FT cells (R70007, ThermoFisher) using Lipofectamine and cell pellets were harvested 24 hours after transfection and snap frozen. Cell pellets were treated with Benzonase (Sigma) in low salt NE1 buffer (20mM HEPES pH7.5, 10mM NaCl, 1mM MgCl_2,_ 0.1% Triton-X100, 10mM β-mercaptoethanol, protease inhibitors) to release chromatin bound proteins and then extracted with 150mM NaCl. Supernatant extracts were mixed with GFP-Trap®_A beads (Chromotek) and after washing beads were boiled in SDS-PAGE sample buffer and released proteins and input extract samples run on 4-15% TGX gradient gels (BioRad) and blotted as above.

Coimmunoprecipitated proteins were detected with the following antibodies: anti-NCoR: rabbit polyclonal A301-145A (Bethyl Laboratories), anti-TBL1X: rabbit polyclonal ab24548 (Abcam), anti-HDAC3: mouse monoclonal 3E11 (Sigma), anti-mSin3a: rabbit polyclonal ab3479 (Abcam), anti-EGFP: mouse monoclonal Living Colors JL-8 (Clontech).

### Methylation-dependent repression assay

WT and mutant mouse *Mecp2* e1 cDNAs were cloned into the expression vector pcDNA3.1(+) IRES GFP (a gift from Kathleen L Collins, Addgene plasmid #51406(52)). The methylation-dependent repression assay was performed as previously described (6) by cotransfecting MeCP2 expression plasmid, Firefly luciferase (Luc) construct pGL2-control (with or without SssI methylation) and Renilla luciferase (Ren) transfection control plasmid into a *Mecp2^−/y^, Mbd2^−/−^* mouse tail fibroblast cell line (6) using Lipofectamine. Luciferase signal was measured using the Dual Luciferase Assay kit (Promega). The ratio Luc/Ren signal was calculated for each well and the mean unmethylated:mean methylated (U/M) value calculated for each genotype. Each combination was repeated in triplicate in each experiment and each MeCP2 genotype was independently transfected at least three times. To check for equal expression of MeCP2 protein, total protein extract from triplicate wells transfected with either unmethylated or methylated luciferase template and each of the MeCP2 expression constructs was western blotted and the level of MeCP2 (normalized to γ-tubulin) determined. Primary antibodies: anti-MeCP2 rabbit monoclonal D4F3 (Cell Signaling Technology), anti-γ-tubulin mouse monoclonal GTU-88 (Sigma).

### Statistical analysis

All statistical analysis was performed using GraphPad Prism (GraphPad Software). Datasets were compared using Student’s t-tests (with or without Welch’s correction for unequal variance, as appropriate), apart from the TBL1X-mCherry colocalisation assay and wire suspension test that were analysed using a non-parametric Mann-Whitney test due to the nature of the data. Rotarod data was tested using two-way repeated measures ANOVA, followed by *post-hoc* testing for significance on each day using the Holm-Sidak method.

### Study approval

All animal experiments were performed under a United Kingdom Home Office project licence (PPL no. 60/4547).

### Data availability

Data from this work are available from the authors.

## Acknowledgements

We would like to thank Alan McClure for animal husbandry and Matthew Lyst and Dirk-Jan Kleinjan for critical reading of the manuscript. A.B is a member of the Simons Initiative for the Developing Brain at the University of Edinburgh.

## Abbreviations

CTD: C-terminal deletion
DAPI: 4’,6-diamidino-2-phenylindole dihydrochloride
EGFP: enhanced green fluorescent protein
ES: cells embryonic stem cells
ExAC: Exome Aggregation Consortium
FACS: fluorescence activated cell sorting
LUHMES: Lund human mesencephalic cells
MBD: methyl-binding domain
NID: NCoR/SMRT-interacting domain
PBS: phosphate-buffered saline
RA: retinoic acid
RTT: Rett syndrome
WT: wild-type

