## Supplementary Materials for "A mutation-led search for novel functional domains in MeCP2"

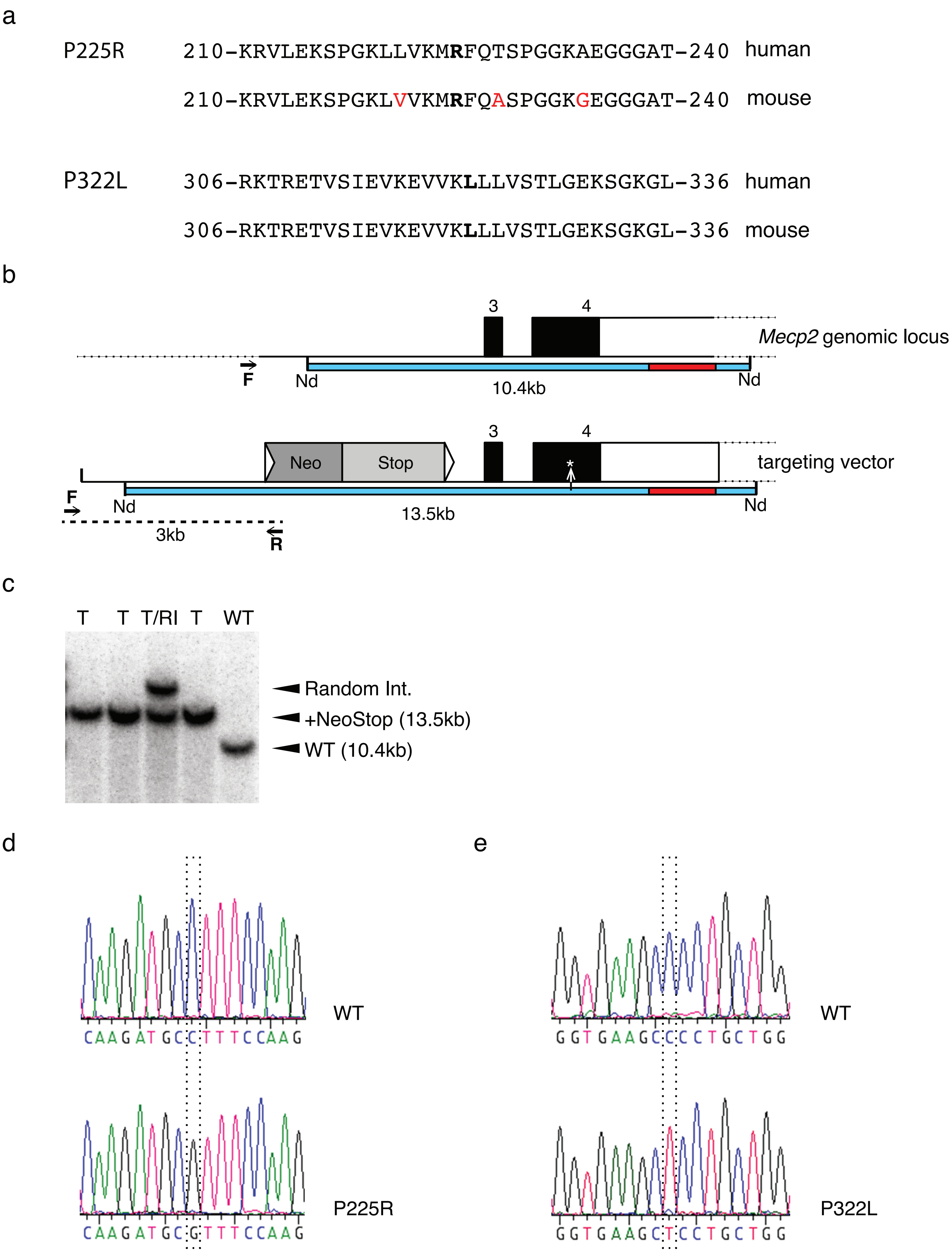


**Supplementary Figure 1**

**Supplementary Figure 1** Production of P225R and P322L knock-in mouse alleles. (**a**) Comparisons of amino acid sequences of human and mouse MeCP2 in the regions flanking RTT missense mutations P225R and P322L. Amino acid differences between human and mouse are in red type and RTT mutations in bold type. (**b**) The mouse *Mecp2* genomic locus and targeting vector showing PCR primers for screening targeted ES clones (black arrows) and genomic NdeI (Nd) fragments used to distinguish correctly targeted clones by Southern blotting. The Southern probe is shown in red and detected fragments in blue. Dotted lines indicate genomic regions outside the targeting vector. (**c**) Southern blot of NdeI digested genomic DNA from WT ES cells (WT), correctly targeted P225R ES cell clones (T), and a targeted clone with an additional randomly integrated copy of the targeting vector (T/RI). (**d**) Sequencing traces confirming presence of mutations P225R and P322L (dashed boxes) in correctly targeted ES cell clones.

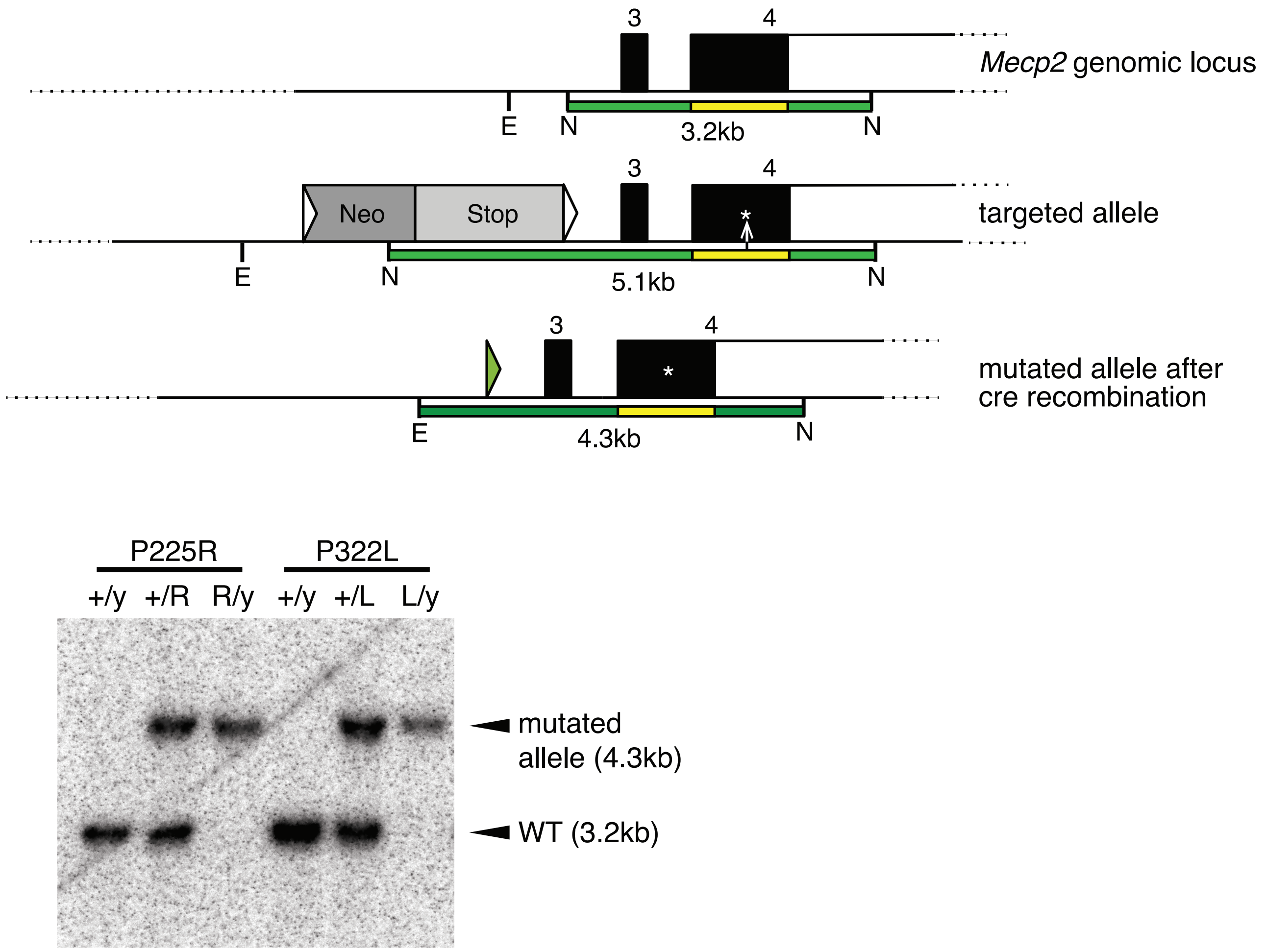


**Supplementary Figure 2** P225R and P322L mouse alleles. (**a**) Structure of the mouse *Mecp2* genomic locus, and the targeted alleles before and after deletion of the NeoStop selection cassette with Cre recombinase. Southern blotting to distinguish between these alleles used an EcoRI (E)/NcoI (N) double digest and *Mecp2* exon 4 probe (yellow). (**b**) Southern blot of genomic DNA from WT and hemizygous mutant male and heterozygous mutant female mice carrying P225R and P322L mutations.

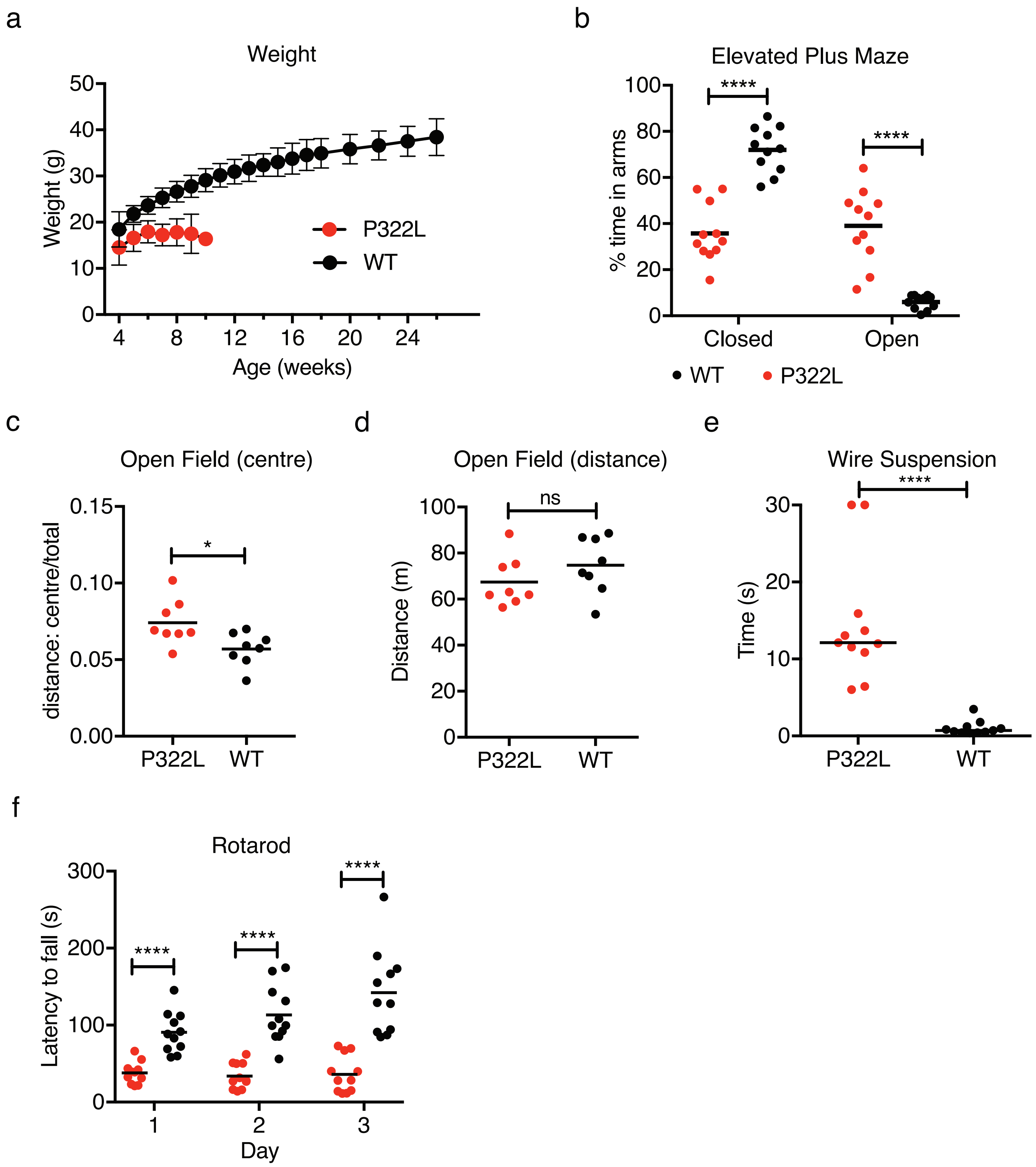


**Supplementary Figure 3** P322L male mice are underweight and show deficits in behavioral tests. (**a**) Mean weight +/- SD for animals shown in Fig. 3a,b (**b**) Elevated plus maze. Percentage of time spent in the closed and open arms is shown as mean (black line) and individual values. Percentage time in open and closed arms for each genotype was compared using two-tailed unpaired t-tests: closed arms P<0.0001 (****), open arms P<0.0001 (****). (continued)

**Supplementary Figure 3 (continued)** (**c**) The ratio of distance travelled in the central zone of the open field arena to total distance travelled shown as mean (black line) and individual values. Two-tailed unpaired t-test test P=0.0188 (*). (**d**) Total distance traveled in the open field arena shown as mean (black line) and individual values. Two-tailed unpaired t-test P=0.2345 (ns). (**e**) Wire suspension test. Time to grasp the wire with the hindlimbs is shown as median (black line) plus individual animal times. Two-tailed Mann-Whitney test P<0.0001 (****). (**f**) Accelerating rotarod test. The latency to fall is shown as mean (black line) and individual values on each day. Two-way ANOVA (repeated measures) [genotype effect, F(1,20) = 45.17, P<0.0001 (****)][time effect, F(2,40)=8.562, P=0.0008 (***)]. Statistical significance was determined between genotypes on each day using the Holm-Sidak method, days 1-3 P<0.0001 (****). (b)(e)(f) P322L n=11, WT n=11. (c)(d) P322L n=8, WT n=8.

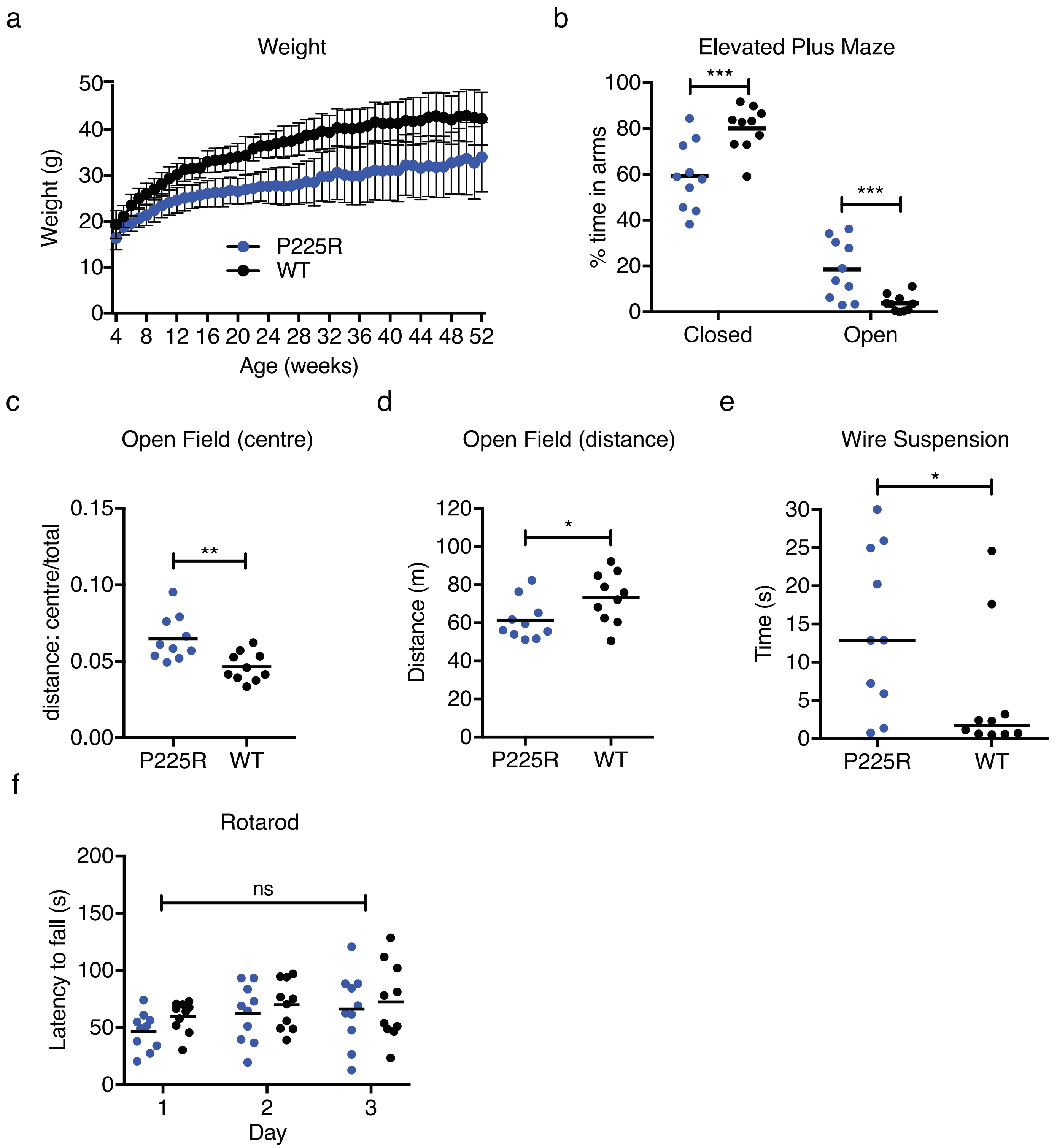


**Supplementary Figure 4** P225R male mice are underweight and show deficits in behavioral tests. (**a**) Mean weight +/- SD for animals shown in Fig. 3e,f. (**b**) Elevated plus maze. Percentage of time spent in the closed and open arms is shown as mean (black line) and individual values. Percentage time in open and closed arms for each genotype was compared using two-tailed unpaired t-tests: closed arms P=0.0016 (***), open arms P=0.0027 (***). (continued)

**Supplementary Figure 4 (continued)** (**c**) The ratio of distance travelled in the central zone of the open field arena to total distance travelled shown as mean (black line) and individual values. Two-tailed unpaired t-test test P=0.0035 (**). (**d**) Total distance traveled in the open field arena shown as mean (black line) and individual values. Two-tailed unpaired t-test P=0.0383 (*). (**e**) Wire suspension test. Time to grasp the wire with the hindlimbs is shown as median (black line) plus individual animal times. Two-tailed Mann-Whitney test P=0.0232 (*). (**f**) Accelerating rotarod test. The latency to fall is shown as mean (black line) and individual values on each day. Two-way ANOVA (repeated measures) [genotype effect, F(1,18) = 0.965, P=0.3390 (ns)]. (b)-(f) P225R n=10, WT n=10.

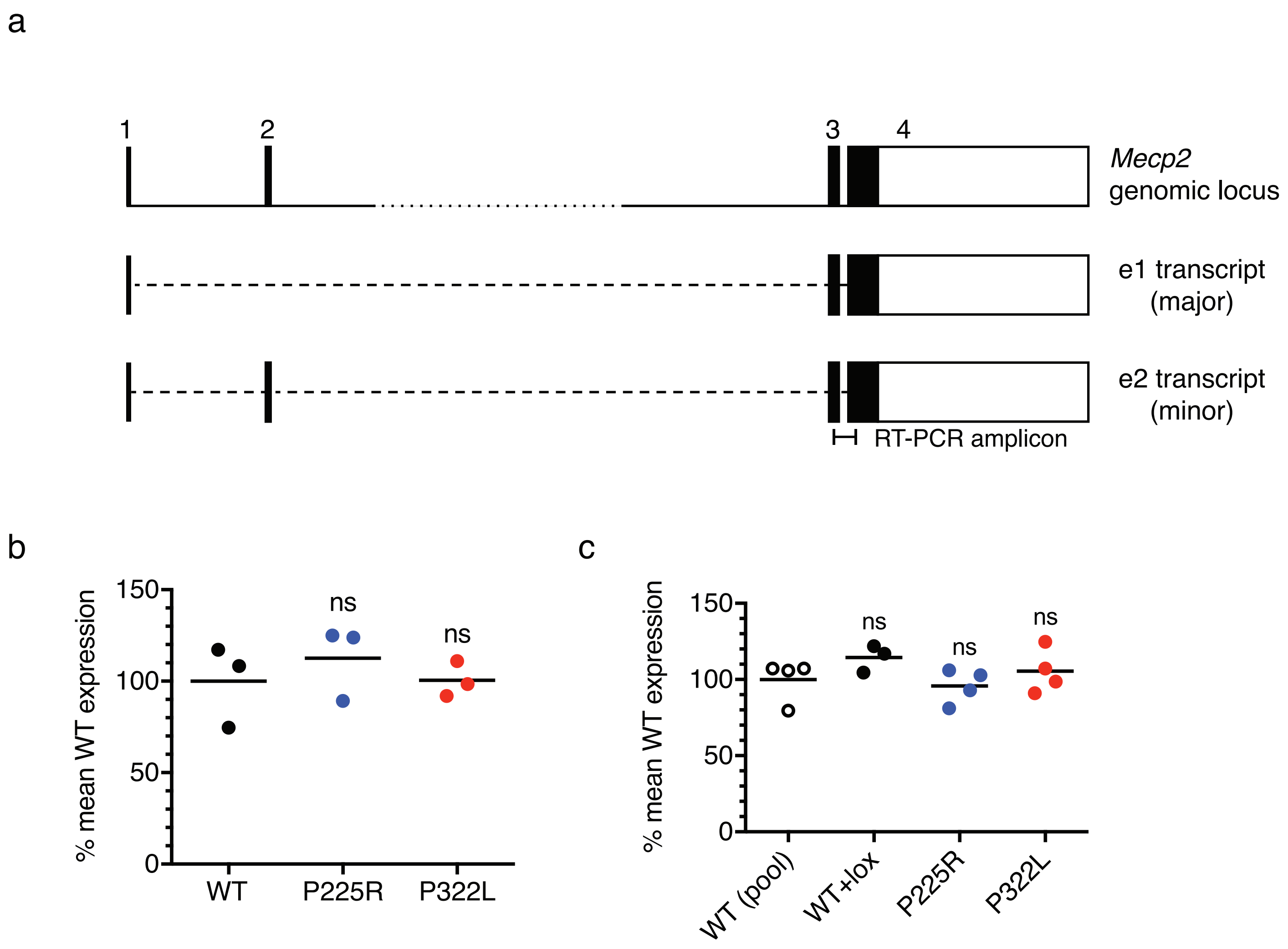


**Supplementary Figure 5** *Mecp2* mRNA levels measured by quantitative RT-PCR. (**a**) *Mecp2* genomic structure and mRNA isoforms e1 and e2. The RT-PCR amplicon detects all *Mecp2* isoforms. (**b**) Quantitative RT-PCR analysis of *Mecp2* mRNA level in whole mouse brain. *Mecp2* expression is relative to *Cyclophilin A* and is normalized to the mean level in WT brain. Expression levels are shown as mean (black line) and values from individual animals. Expression levels were compared to WT using two-tailed unpaired t-tests (all n=3). P225R P=0.5098 (ns), P322L P=0.9751 (ns).(**c**) Quantitative RT-PCR analysis of *Mecp2* mRNA level in in vitro differentiated neurons (7 days of differentiation). *Mecp2* expression is relative to *Cyclophilin A* and is normalized to the mean level in WT neurons. Expression levels are shown as mean (black line) and individual values from independent differentiations. Comparison to WT (n=4) using two-tailed unpaired t-tests: WT+lox (n=3) P=0.1513 (ns), P225R (n=4) P=0.6484 (ns), P322L (n=4) P=0.3570 (ns).

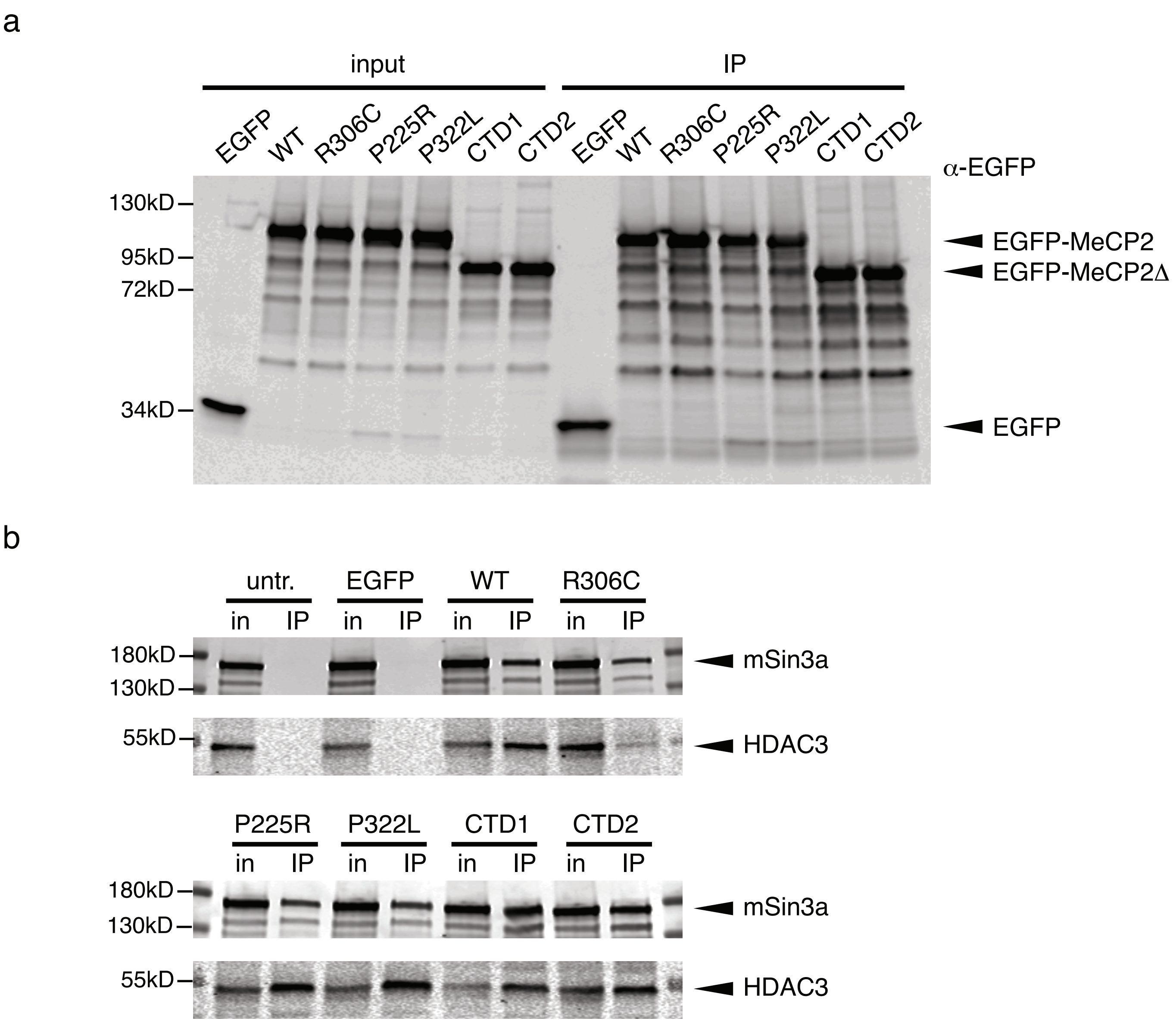


**Supplementary Figure 6** Immunoprecipitation of transfected EGFP-MeCP2 from HEK 293 cells. (**a**) Western blot with input and immunoprecipitated samples from Fig. 4b probed with an antibody against EGFP to show equivalent expression and immunoprecipitation of all EGFP-MeCP2 genotypes. (**b**) Immunoprecipitation of EGFP-MeCP2 from transfected HEK 293 cells with GFP-Trap beads. The corepressor mSin3a, in addition to NCoR/SMRT component HDAC3 is detected in western blots of input and immunoprecipitated samples.

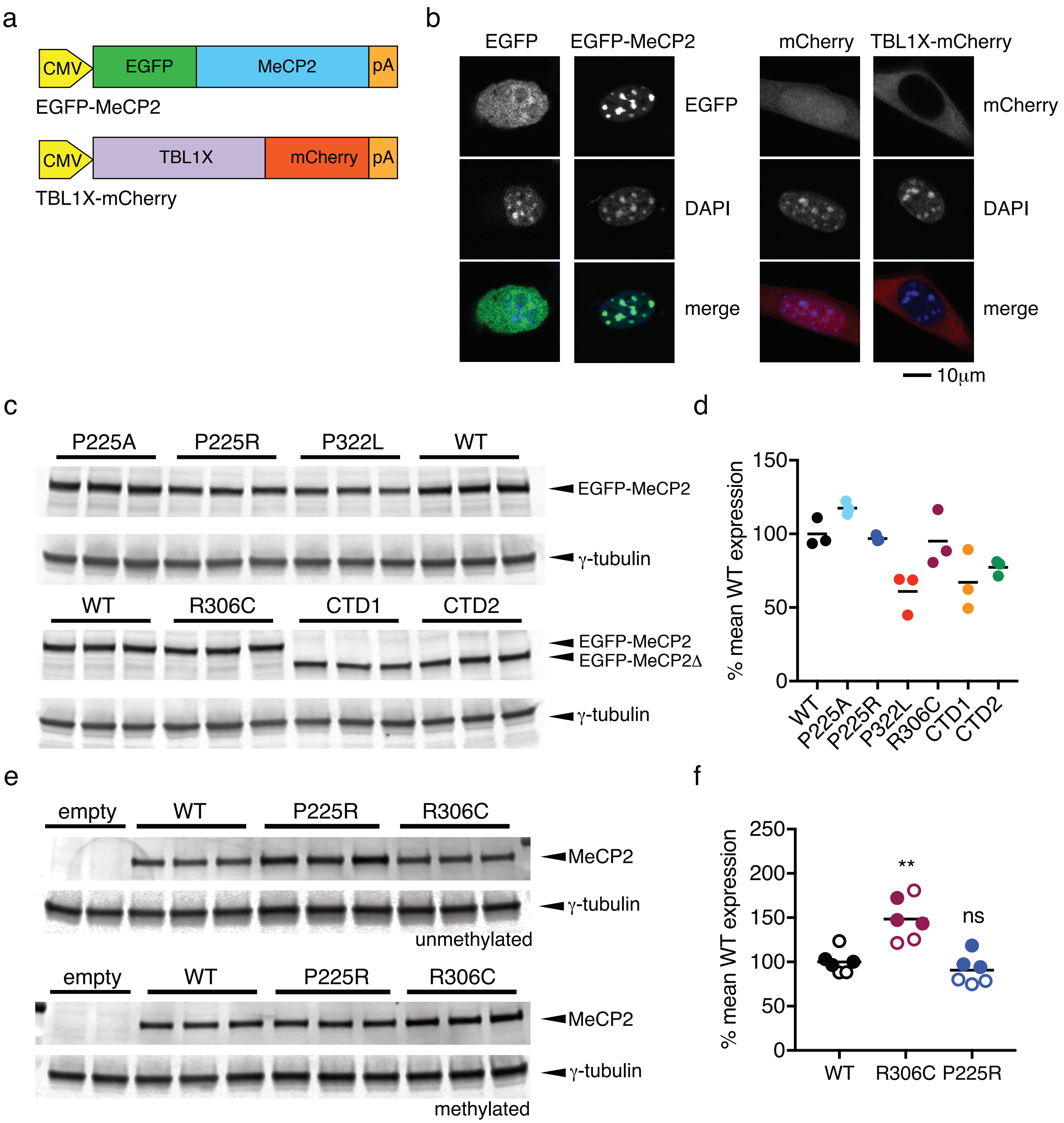


**Supplementary Figure 7** TBL1X-mCherry recruitment assay and luciferase methylation-dependent repression assay. (**a**) Schematic illustration of EGFP-tagged MeCP2 and mCherry-tagged TBL1X expression constructs. (**b**) EGFP fluorescence is taken to heterochromatic foci in the nucleus when fused to MeCP2. mCherry fluorescence is excluded from the nucleus when fused to TBL1X. (continued)

**Supplementary Figure 7 (continued)** (**c**) Western blot to show that MeCP2 WT and mutant proteins are expressed at equivalent levels in the TBL1X-mCherry recruitment assay shown in Fig. 4d. Each lane shows protein from one well of NIH 3T3 cells transfected with EGFP-MeCP2 and TBL1X-mCherry (n=3 for each genotype). Western blot probed with an antibody that recognizes both full-length and truncated MeCP2, and γ-tubulin as a loading control. (**d**) Quantification of (c). MeCP2/γ-tubulin expression normalized to mean WT EGFP-MeCP2 level and plotted as mean (black line) and individual values. (**e**) Western blot to show that MeCP2 WT and mutant proteins are expressed at equivalent levels in the methylation-dependent transcriptional repression assay shown in Fig. 4f. Each lane shows protein from one well of Mecp2-/y, Mbd2-/- mouse tail fibroblasts transfected with SssI-methylated (upper panels) or unmethylated (lower panels) pGL2-Control plasmid and an MeCP2 expression construct (n=3 for each combination). Western blot probed with an antibody against MeCP2, and γ-tubulin as a loading control. (**f**) Quantification of (e). MeCP2/γ-tubulin expression normalized to mean WT level and plotted as mean (black line) and individual values. Open circles – unmethylated template, filled circles – methylated template. Comparison to WT using two-tailed unpaired t-tests: R306C P=0.0015 (**), P225R P=0.3001 (ns). All genotypes n=6.

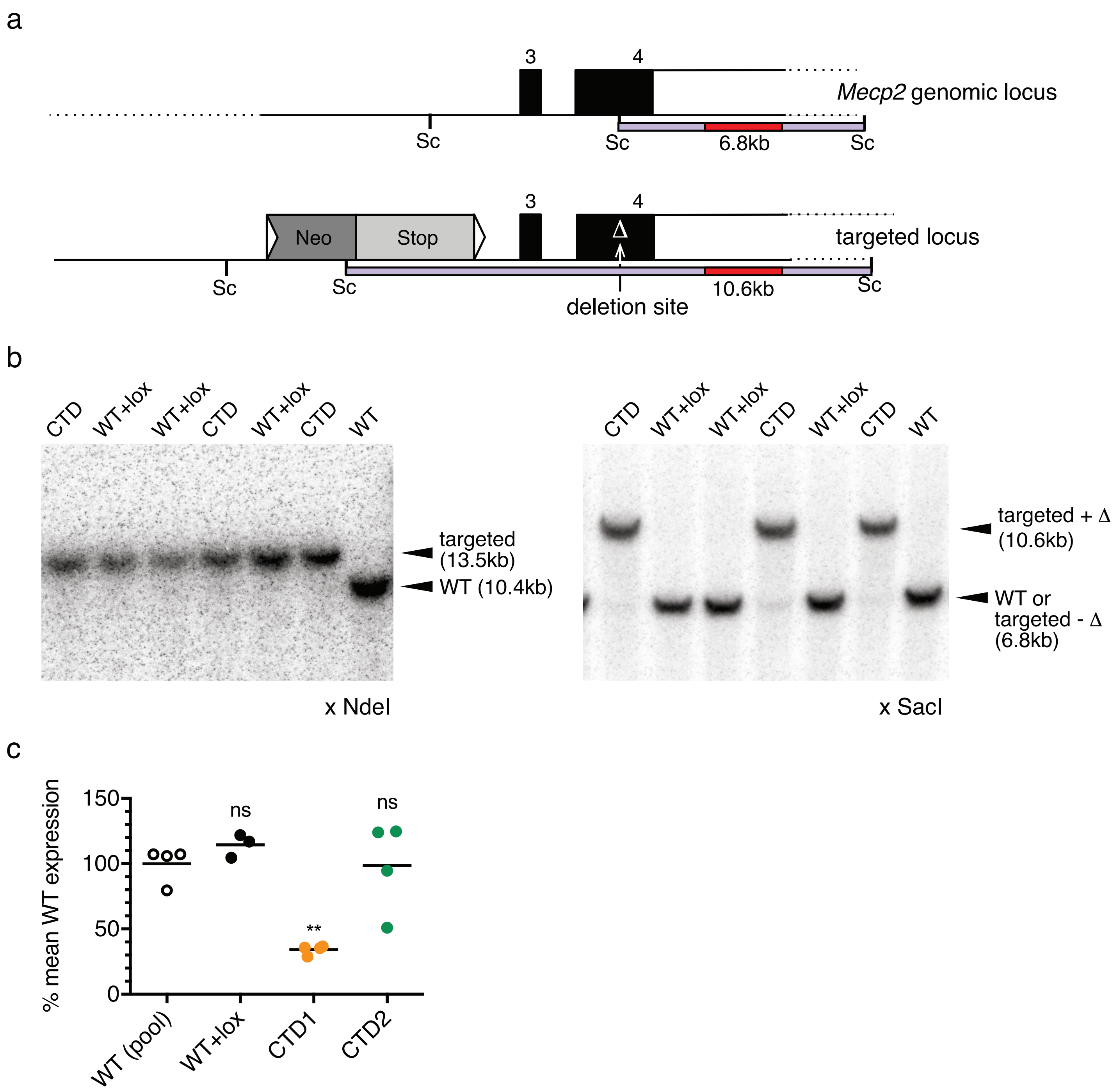


**Supplementary Figure 8** Production of CTD1 and CTD2 knock-in alleles in mouse ES cells. (**a**) The mouse *Mecp2* genomic locus and targeted locus showing genomic SacI (Sc) fragments used to distinguish correctly targeted clones by Southern blotting. The Southern probe is shown in red and detected fragments in lilac. Dotted lines indicate genomic regions outside the targeting vector. (**b**) Southern blots of NdeI (see Supp. Fig. 1b) and SacI digested genomic DNA from WT ES cells (WT), correctly targeted ES cell clones with the CTD1 deletion (CTD) and targeted clones which have been targeted with the selection cassette but do not have the deletion (WT+lox). (continued)

**Supplementary Figure 8 (continued)** (**c**) Quantitative RT-PCR analysis of *Mecp2* mRNA level in *in vitro* differentiated neurons (7 days of differentiation). *Mecp2* expression is relative to *Cyclophilin A* and is normalized to the mean level in WT neurons. Expression levels were compared to WT (n=4) using two-tailed unpaired t-tests with Welch's correction for unequal variance. WT+lox (n=3) P=0.1513 (ns), CTD1 (n=4) P=0.0016 (***), CTD2 (n=4) P=0.7851 (ns).

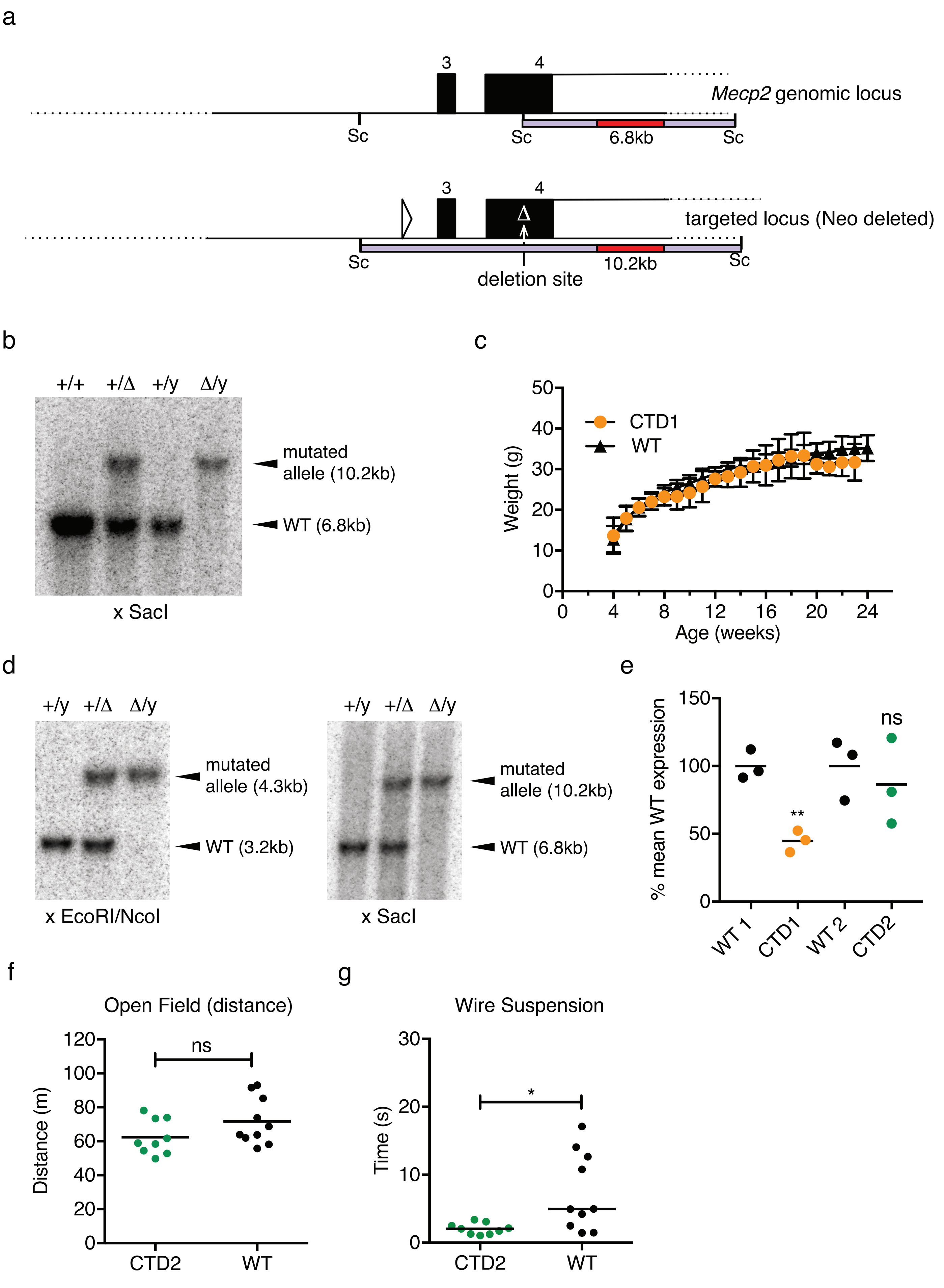


**Supplementary Figure 9**

**Supplementary Figure 9** CTD1 and CTD2 mouse alleles. (**a**) Structure of the mouse *Mecp2* genomic locus and the targeted allele after deletion of the NeoStop selection cassette with Cre recombinase (equivalent to the CTD1 allele generated by CRISPR/Cas9 cutting and oligonucleotide repair in oocytes). Southern blotting to distinguish between these alleles used an EcoRI/NcoI double digest as shown in Supp. Fig. 3a or a SacI (Sc) digest as shown in Supp. Fig 7a. (**b**) Southern blot of SacI-digested genomic DNA from WT and hemizygous mutant male and WT and heterozygous mutant female mice carrying the CTD1 mutation. (**c**) Mean weight +/- SD for animals shown in Fig. 6c,d. (**d**) Southern blots of EcoRI/NcoI and SacI-digested genomic DNA from WT and hemizygous mutant male and heterozygous mutant female mice carrying the CTD2 mutation. (**e**) Quantitative RT-PCR analysis of *Mecp2* mRNA level in whole mouse brain. *Mecp2* expression is relative to *Cyclophilin A* and is normalized to the mean level in WT littermate brains. Data shown as mean (black line) and individual animals. Expression levels were compared to WT littermate controls using two-tailed unpaired t-tests (all n=3). CTD1 P=0.0029 (**), CTD2 P=0.5818 (ns). (**f**) Total distance traveled in the open field arena shown as mean (black line) and individual values. Two-tailed unpaired t-test P=0.1200 (ns). (**g**) Wire suspension test for CTD2 and WT males. Time to grasp the wire with the hindlimbs is shown as median (black line) plus individual animal times. Two-tailed Mann-Whitney test P=0.0101(*). (f),(g) CTD2 n=9, WT n=10.

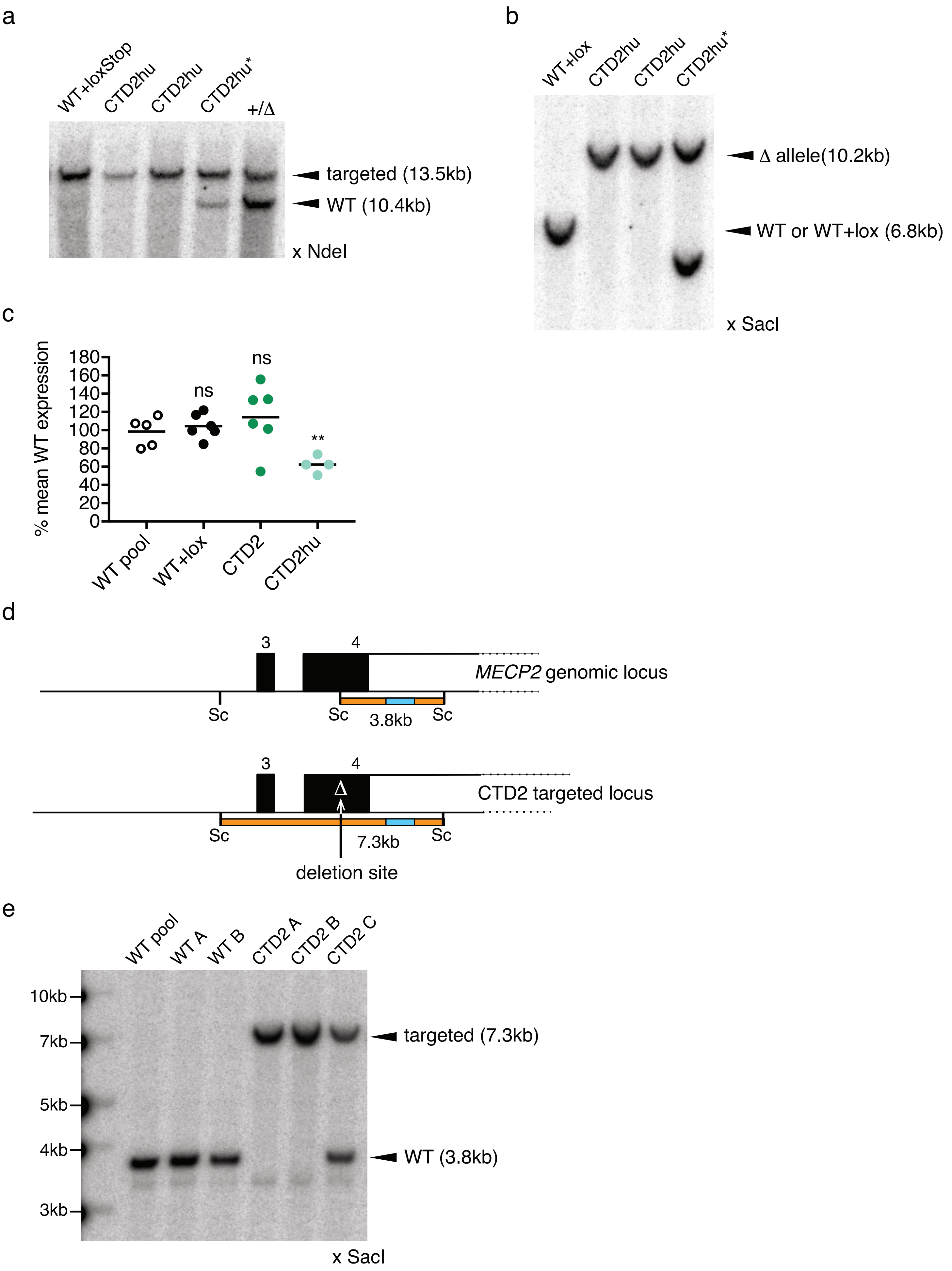


**Supplementary Figure 10**

**Supplementary Figure 10** Production of humanised mouse and human CTD2 alleles. (**a**) Southern blot of NdeI digested genomic DNA from correctly targeted ES cell clones with the CTD2hu deletion and a clone that has been targeted with the selection cassette but does not have the deletion (WT+lox). CTD2hu* has been targeted correctly but also contains an additional band so was not used. +/Δ is genomic DNA from a heterozygous CTD2 female mouse that carries both WT and CTD alleles. (**b**) Southern blot of SacI digested genomic DNA from correctly targeted ES cell clones with the CTD2hu deletion as in (a). (**c**) Quantitative RT-PCR analysis of *Mecp2* mRNA level in in vitro differentiated neurons (7 days of differentiation). *Mecp2* expression is relative to *Cyclophilin A* and is normalized to the mean level in WT neurons. Data is plotted as mean (black line) and individual differentiations. Expression levels were compared to WT (n=5) using two-tailed unpaired t-tests. WT+lox (n=6) P=0.5236 (ns), CTD2 (n=6) P=0.3817 (ns), CTD2hu (n=4) P=0.0052 (**). (**d**) Structure of the human *MECP2* genomic locus and the targeted CTD2 allele created by CRISPR/Cas9 cutting and oligonucleotide repair in LUHMES cells. Southern blotting to distinguish between these alleles used a SacI digest (Sc) and a probe containing part of the 3'UTR (blue). Resulting fragments are shown in orange. (**e**) Southern blot of SacI digested genomic DNA from LUHMES CTD2 knock-in clones and unmodified controls. CTD2 A and B carry homozygous CTD2 knock-ins. CTD2 C is heterozygous with one active CTD2 allele and one inactive WT allele.
